# A new form of actin assembly around the *Shigella*-containing vacuole regulates its intracellular niche

**DOI:** 10.1101/2020.03.12.988097

**Authors:** Sonja Kühn, John Bergqvist, Laura Barrio, Stephanie Lebreton, Chiara Zurzolo, Jost Enninga

## Abstract

The enteroinvasive bacterium *Shigella flexneri* forces its uptake into non-phagocytic host cells through the translocation of T3SS effectors that subvert the actin cytoskeleton. Here, we report *de novo* actin polymerization after cellular entry around the bacterial containing vacuole (BCV) leading to the formation of a dynamic actin cocoon. This cocoon is thicker than any described cellular actin structure and functions as a gatekeeper for the cytosolic access of the pathogen. Host Cdc42, Toca-1, N-WASP, WIP, the Arp2/3 complex, cortactin, coronin, and cofilin are recruited to the actin cocoon. They are subverted by T3SS effectors, such as IpgD, IpgB1, and IcsB. IcsB immobilizes components of the actin polymerization machinery at the BCV. This represents a novel microbial subversion strategy through localized entrapment of host actin regulators causing massive actin assembly. We propose that the cocoon protects *Shigella*’s niche from canonical maturation or host recognition.

## INTRODUCTION

Bacterial pathogens have evolved sophisticated ways to drive infection and to establish their intracellular niches for survival and replication. Especially the cellular actin cytoskeleton is extensively hijacked for bacterial purposes. This cytoskeletal meshwork is controlled by a complex network of actin-binding proteins (ABPs), which nucleate new actin filaments (F-actin) from actin monomers (G-actin), or elongate, maintain and disassemble existing ones. ABPs are spatiotemporally localized and regulated by Rho GTPases, phospholipids, post-translational modifications, or membrane-bound scaffold (Le Clainche and Carlier, 2008, Rottner et al., 2017, Pollard, 2016). The main F-actin nucleating factor is the Arp2/3 complex (Machesky et al., 1994) (Mullins et al., 1998), which generates branched actin meshworks in membrane ruffles and lamellipodia, at phagosomes and intracellular vesicles. Yet efficient F-actin nucleation requires additional nucleation-promoting factors (NPFs) like N-WASP. N-WASP is itself activated by several factors like PI(4,5)P_2_, the F-BAR scaffold Toca-1, and the Rho GTPase Cdc42 (Ho et al., 2004, Rohatgi et al., 1999, Rottner et al., 2017). Rho GTPases are central regulators of the actin cytoskeleton and switch between an inactive, GDP-bound and an active, GTP-bound state (Vetter and Wittinghofer, 2001). This is controlled by guanine nucleotide-exchange factors (GEFs) promoting GDP dissociation, GTPase-activating proteins (GAPs), and guanine nucleotide dissociation inhibitors (GDIs) (Etienne-Manneville and Hall, 2002) (Cherfils and Zeghouf, 2013). In the active state, Rho GTPases interact with effector proteins for cell signaling and to regulate the actin cytoskeleton. Remarkably, bacterial pathogens like the Gram-negative, enteroinvasive bacterium *Shigella flexneri* (hereafter *Shigella*) do not directly modify actin (Kühn and Mannherz, 2017). Instead, *Shigella* modulates the recruitment and the activation of actin regulators by subverting upstream Rho GTPases, kinases, and phospholipid signaling (Schnupf and Sansonetti, 2019) (Schroeder and Hilbi, 2008) (Valencia-Gallardo et al., 2015).

*Shigella* is the causative agent of bacterial dysentery and an important model for intracellular pathogenesis (Schnupf and Sansonetti, 2019). It forces its uptake into non-phagocytic epithelial cells through the translocation of type 3 secretion system (T3SS) effectors. These proteins target the host actin cytoskeleton and endomembrane trafficking to induce cellular entry and to establish an intracellular replicative niche. For cellular entry, thin membrane protrusions make the first contact with bacteria, followed by the initiation of massive actin rearrangements enclosing the entering *Shigella* (Schroeder and Hilbi, 2008) (Valencia-Gallardo et al., 2015) (Cossart and Sansonetti, 2004) (Romero et al., 2012). After cellular uptake in a tight bacterium-containing vacuole (BCV) (Weiner et al., 2016), *Shigella* induces its rapid escape for replication into the host cytosol. There, it recruits the host actin nucleation machinery to one of its poles by its virulence factor IcsA to spread from cell-to-cell (Suzuki et al., 1998, Egile et al., 1999, Gouin et al., 1999). Parallel to its uptake, *Shigella* induces the formation of infection-associated macropinosomes (IAMs). These IAMs accumulate at the entry site and surround the BCV. They form membrane-membrane contacts with the ruptured BCV and their presence correlates with efficient rupture (Mellouk et al., 2014, Weiner et al., 2016).

We have recently discovered the formation of a hitherto undescribed actin cytoskeleton structure that assembles around vacuolar *Shigella* (Ehsani et al., 2012, Mellouk et al., 2014, Weiner et al., 2016). Here, we performed its in-depth characterization coining it as “actin cocoon”. We found that this cocoon is thicker than any other cellular actin structure and only assembles after bacterial uptake. We identified the process underlying its formation, namely the involved bacterial T3SS effectors and a subverted host pathway for actin rearrangements. Finally, we demonstrate that interfering with cocoon formation and disassembly affect *Shigella*’s capacity to invade the host cytosol.

## RESULTS

### The actin cocoon assembles *in situ* after cellular entry around *Shigella*-containing phagosomes and its disassembly precedes cytosolic escape

Actin-GFP transfected HeLa cells were imaged during early infection steps of wild type, dsRed-expressing *Shigella* at high spatiotemporal resolution (Figure 1A,B). After two hours, almost all cells were infected with no further primary infection, and membrane ruffling was shut down. Live imaging (Figure 1A,B) and fixed experiments (Figure S1A,B) revealed the *in situ* assembly of a thick actin coat-like structure after pathogen entry as indicated by a massive increase in fluorescence intensity around the bacterium-containing vacuole (BCV). This “actin cocoon”-termed structure was distinct from cortical actin and polymerized *de novo* at the surface of the entire vacuolar membrane. After a fast nucleation phase of 1-3 min, the actin cocoon was maintained until its final disassembly, which was immediately followed by BCV membrane rupture (Figure 1A-C). All observed actin rearrangements took place in the time span after entry site formation and before the cytosolic spread of *Shigella*. Phalloidin-staining of endogenous actin in fixed experiments revealed the presence of F-actin in the cocoon of *Shigella* invading HeLa or Caco-2 cells (Figure S1A,B).

**Figure 1.**
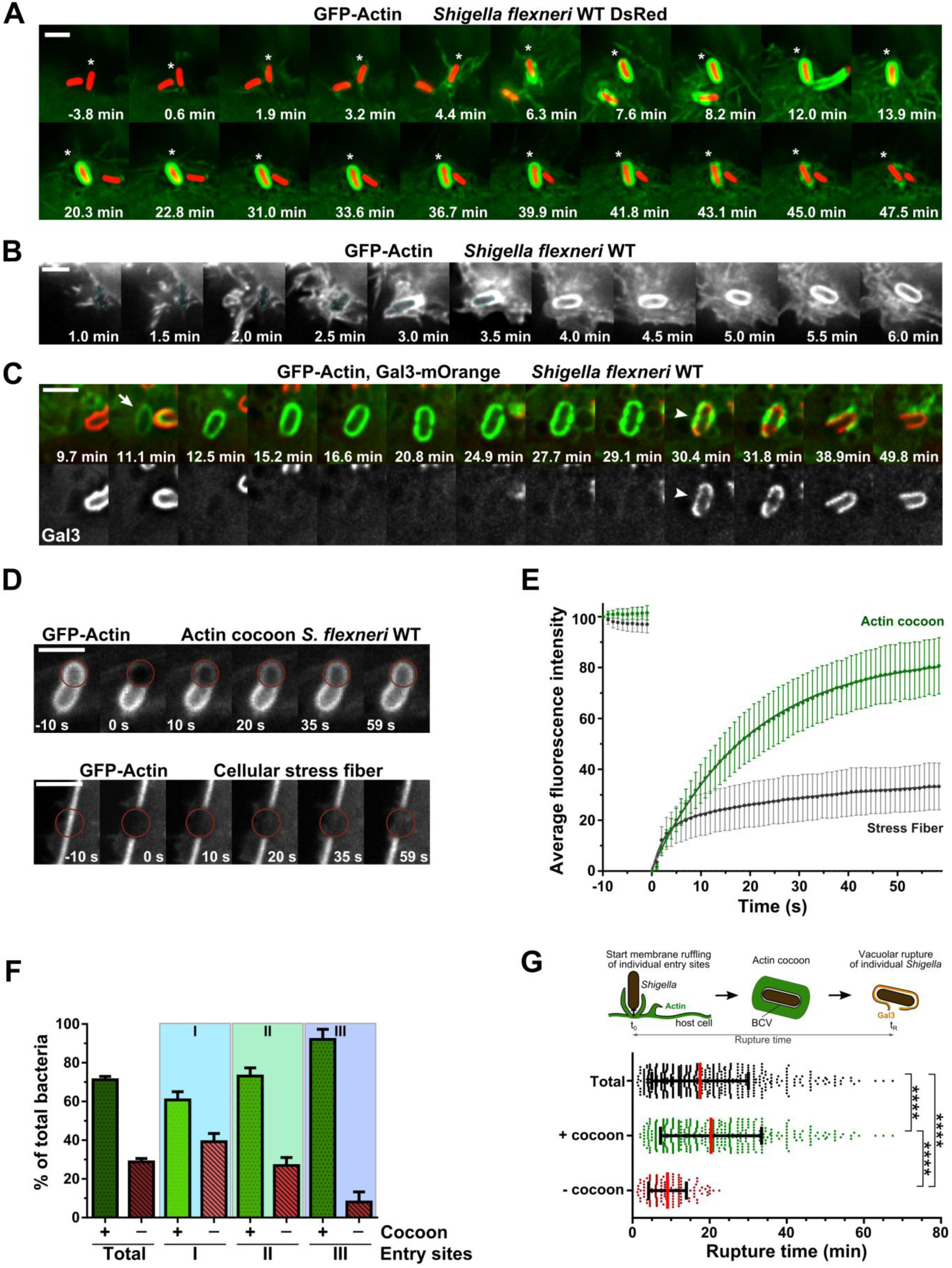
The dynamic actin cocoon polymerizes *de novo* after cellular entry around *Shigella*’s BCV and disassembles before vacuolar escape. (**A-B**) Real time-monitoring of the thick actin cocoon. HeLa cells expressing actin-GFP (green) were infected with *Shigella* WT DsRed (red) (A, *: bacterium with cocoon) or *Shigella* WT (B, blue dashed line: bacterium). t=0 min: onset of entry site formation. (**C**) The actin cocoon needs to at least partially disassemble before rupture. Time-lapse of *Shigella WT* infecting HeLa cells expressing actin-GFP (green) and galectin-3-mOrange (red, Gal3). Arrow: newly formed cocoon, arrow head: moment of vacuolar rupture. (**D-E**) FRAP experiments reveal constant F-actin turnover of the cocoon and different dynamics compared to stress fibers (actin cocoon: n=3, N=40, green; stress fibers: n=5, N=75, grey). Plotted are mean values ± SD and curve fit. (**F-G**) Actin cocoon formation depends on the time point of bacterial infection and pre-infection. Depicted are percentages of bacteria that successfully escaped into the host cytosol and previously assembled an actin cocoon (+) or not (-) (F). On average, three entry sites formed per cell (Figure S2) and 71.2±1.75% of invading bacteria (n=4, N=631) had a cocoon. All late invaders assembled a cocoon before vacuolar escape (G). Mean values ± SD of individual experiments (F), or rupture time points of single invaders (G) are plotted. Mann-Whitney test with p<0.05 as significant: (****p<0.0001). Scale bars: 3 μm. (See Figure S1, S2, S3)

To monitor the precise time point of vacuolar rupture in correlation to actin cocoon formation, we used fluorescently-labeled galectin-3 as marker. At the moment of vacuolar membrane damage, galectin-3 molecules get recruited to β-galactosides at the inner leaflet of the phagosomal membrane. We never observed vacuolar rupture by bacteria residing in an intact cocoon. At least a partial disassembly of the actin cocoon always preceded the directly following vacuolar rupture (Figure 1C). We conclude that cocoon disassembly is tightly linked with bacterial release into the host cytosol.

### The actin cocoon is dynamically re-assembled during its lifetime and its formation depends on the time point of infection

Next, we performed fluorescence recovery after photobleaching (FRAP) measurements to compare the spatiotemporal dynamics of the actin cocoon with cellular stress fibers (Figure 1D,E). We anticipated fluorescence recovery either from F-actin treadmilling and *de novo* polymerization, or to a lesser extent from free diffusion of cytoplasmic G-actin (a very fast saturated process). Strikingly, actin filaments of the cocoon had a high turnover rate with a mobile fraction (F_m_) of 84.6±0.77% and a halftime of fluorescence recovery (t_1/2_) of 13.8±0.79 s (Figure 1E). This revealed constant incorporation of new, unbleached G-actin and thus ongoing F-actin polymerization throughout the cocoon until its final disassembly. The same was observed by imaging with constant partial disassembly and re-assembly of the cocoon (Figure S1C). Actin turnover in cellular stress fibers occurs through the incorporation of new actin monomers or filaments. We found the turnover of the cocoon to be clearly different compared to cellular stress fibers (F_m_=36.8±2.14%) (Figure 1E). This points to constant *de novo* F-actin nucleation or elongation during its entire lifetime.

To quantify *Shigella* infection with regards to actin cocoon formation and vacuolar rupture, we monitored the successive infection steps of individual bacteria (Figure S2). Strikingly, in total 71.2±1.75% of all *Shigella* assembled a dynamic actin cocoon before cytosolic escape (Figure 1F). *Shigella*’s probability for cocoon formation was pronounced in cells which had already been infected. This occurred with the same tendency, either if several bacteria entered via the same or via different entry sites (Figure 1F, Figure S2D). We also quantified the time span that individual *Shigella* required after the start of entry site formation (initial cortical actin rearrangements) to escape into the host cytosol (initial galectin-3 recruitment). We found that all late invading bacteria polymerized an actin cocoon as shown by a significant delay in rupture time (-cocoon: 9.48±5.63 min, +cocoon: 20.37±13.13 min, p<0.0001, Figure 1G). Increasing the multiplicity of infection (MOI>15) increased the infection efficiency with more bacteria entering via the same entry site, but not the probability of cocoon formation and rupture time (Figure S3). Taken together, we conclude (i) that cocoon assembly depends on the phagocytic load as well as the order of infection, and (ii) that all late invading bacteria assemble an actin cocoon.

### The *Shigella*-specific actin cocoon represents a unique structure

To better understand the nature of actin cocoons, we compared them with well-characterized host actin cytoskeletal structures. We examined by quantitative fluorescence microscopy structures that are either very dynamic, like the actin meshwork at lamellipodial tips, or very thick, like cellular stress fibers of F-actin bundles (Figure 2A). We measured the fluorescence intensities of the thickest stress fibers per infected cell and normalized each individual measurement against the average stress fiber intensity (Supplemental Information). Our analysis revealed that actin cocoons are much denser than any other actin structure with an on average 7-fold higher fluorescence intensity compared to stress fibers (7.02±1.25, p<0.0001), and 11.5-fold compared to the lamellipodial tip (0.61±0.10, p<0.0001) (Figure 2A). In line with this, our previously published CLEM data sets of three actin cocoons (Weiner et al., 2016) exhibited a maximum thickness of 350 nm. Additionally, the cocoon differed from short-lasting actin rearrangements around *Shigella*-induced macropinosomes, which did not differ significantly in fluorescence intensity compared to stress fibers (1.30±0.57, Figure 2A), resembling the actin flashing phenomenon (Yam and Theriot, 2004) (Liebl and Griffiths, 2009) (see below).

**Figure 2.**
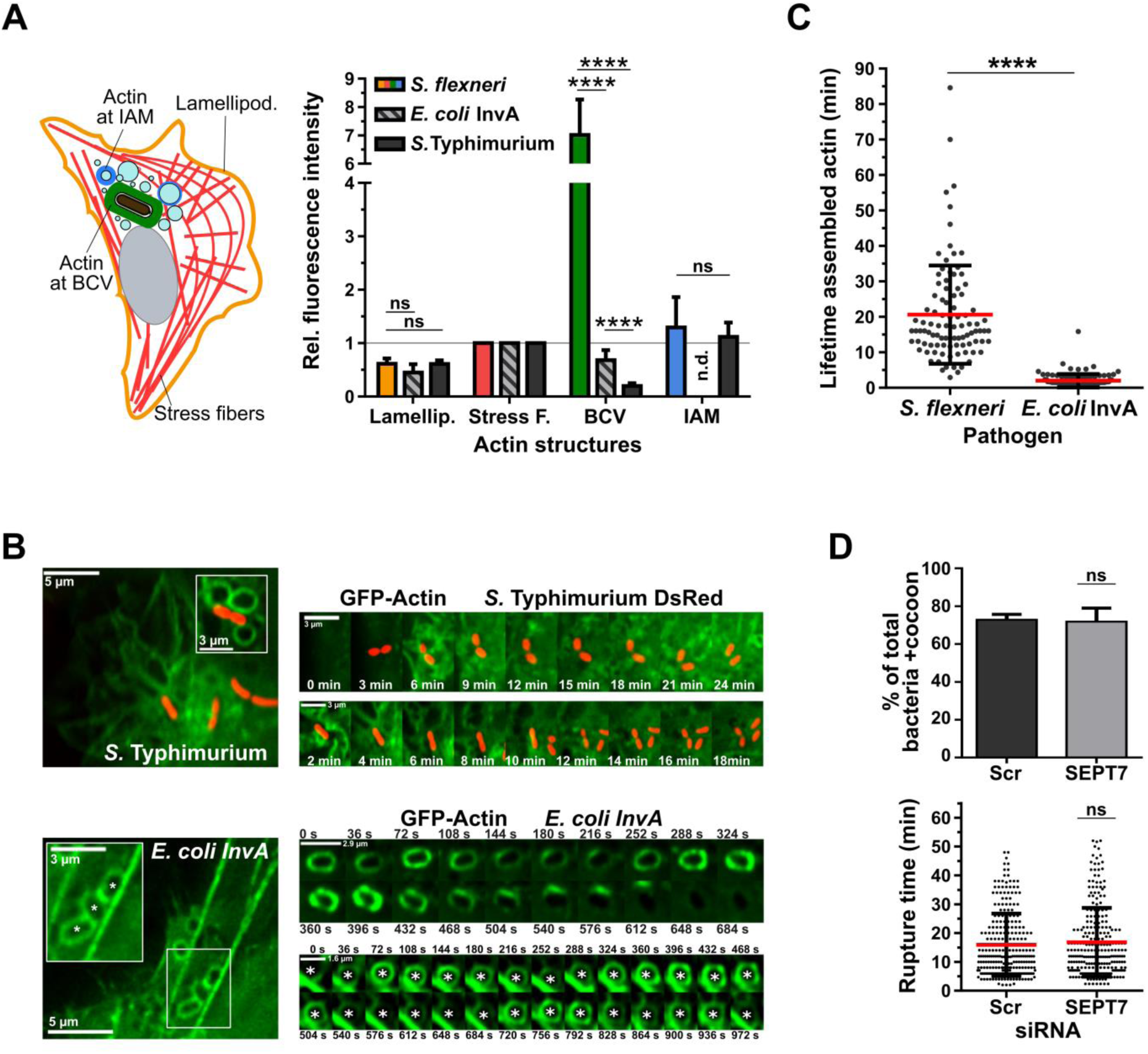
The actin cocoon is a new *Shigella*-specific cytoskeletal structure. (**A**) The actin cocoon is thicker than any cellular actin structure and *Shigella-*specific. Left: Scheme of investigated actin structures with stress fibers (Stress F., red), lamellipodial tip (Lamellip., orange), actin at BCVs (green) and at IAMs (blue). Right: Quantification of relative fluorescence intensities. All measurements were normalized per cell to the average fluorescence intensity of the thickest stress fibers (*S. flexneri* WT: n=4, N=250; *E. coli* InvA: n=3, N=236; *S.* Typhimurium WT: n=3, N=322). (**B**) Time-lapse imaging examples of (A). Top: Invading *S.* Typhimurium did not assemble an actin cocoon. Bottom: Cyclic actin flashing around *E. coli* InvA phagosomes (*) in comparison with cellular stress fibers. (**C**) The actin cocoon lifetime of *Shigella* WT (N=95) differs from actin flashing by *E. coli* InvA (N=103) and is 10-times longer (20.64±13.87 min). (**D**) The actin cocoon assembles independently of septin-7-containing filaments as seen with knockdown of SEPT7 by siRNA. (Scr: scramble, control, N=264. SEPT7: N=250, KD 91.7%, see Figure S4). Indicated are mean values ± SD (n=3). Statistical significance: p<0.05 as significant (****p<0.0001, ns: not significant).

Next, we deciphered if the actin cocoon is specific for *Shigella* or a general mechanism occurring during cell entry. Therefore, we compared it with actin rearrangements during early infections of the closely related bacterium *Salmonella enterica* serovar Typhimurium (*S.* Typhimurium). Like *Shigella, Salmonellae* induce massive membrane ruffling and macropinocytosis during cellular entry by T3SS effector proteins. To the contrary, *S.* Typhimurium pursue mainly an intravacuolar lifestyle for bacterial replication (Santos and Enninga, 2016). To decipher actin polymerization around entering *Salmonella*, we followed bacteria-induced actin rearrangements during the first 2 hrs post infection. (Figure 2A,B). We barely detected any around *Salmonella*-containing vacuoles (SCV) (35-fold less, 0.20±0.05) (Figure 2A,B). Instead, actin occurred as small, intense dots around the SCV, probably derived from recycling and fusion events for vacuolar maturation. We concluded that *Salmonella* does not assemble an actin coat-like structure during early invasion steps.

Further, we examined the host cell contribution for cocoon assembly. Previous work of several groups described short-lasting, repeated cycles of actin polymerization and depolymerization around fully internalized phagosomes (Yam and Theriot, 2004) (Liebl and Griffiths, 2009). These ‘actin flashing’ termed actin rearrangements were identified as downstream consequences of several cellular entry mechanisms independent of cell type and cargo in cells with high phagocytic load (Liebl and Griffiths, 2009). To compare actin flashes with actin cocoons, we infected HeLa cells with *Escherichia coli* expressing *Yersinia pseudotuberculosis* invasin A (*E. coli* InvA) as model for canonical phagocytosis. Invasin A binds integrins and is sufficient to transfer *Yersinia*’s zippering entry to *E. coli* (Isberg et al., 1987). We observed successive waves of actin flashing around internalized *E. coli* InvA with 32% lower fluorescence intensity compared to stress fibers (0.68±0.19, p=0.0056). In line with previous studies (Yam and Theriot, 2004) (Liebl and Griffiths, 2009), the duration was 2.0±1.9 min per cycle (Figure 2A-C). Thus, actin cocoons were distinct from the ‘actin flashing’ phenomenon in several major points: First, the fluorescence intensity of *Shigella*’s cocoons was on average 10-fold higher (Figure 2A). Second, the cocoon had a more than 10-times longer lifespan than one actin flash (Figure 2C). Third, we did not identify any consecutive cycles of actin assembly at *Shigella*’s BCV. Cocoon disassembly was always followed by immediate cytosolic escape, while *E. coli* InvA phagosomes never ruptured. In conclusion, the actin cocoon around vacuolar *Shigella* clearly differs from transient actin structures around endocytic compartments.

Finally, we examined if other cytoskeletal systems like septin filaments (Kinoshita et al., 2002) (Mavrakis et al., 2014) contribute to the assembly of this new actin structure. During late infection steps, cytosolic *Shigella* are trapped in septin-actin cages to restrict bacterial proliferation (Mostowy and Cossart, 2012, Mostowy et al., 2011, Mostowy et al., 2010). We performed knockdown of SEPT7 by RNA interference, a common technique for the efficient inhibition of septin filament formation (Sirianni et al., 2016). This did neither prevent cocoon formation, nor delay *Shigella*’s cytosolic escape (Figure 2D, Figure S4A). Therefore, the actin cocoon around vacuolar *Shigella* forms independently from septins.

### The host Arp2/3 complex-dependent actin nucleation machinery is recruited to the BCV

In order to identify host factors that initiate actin cocoon formation, we performed an inhibitor screen targeting host actin regulators. A general complication in examining intracellular, pathogen-induced actin cytoskeleton subversions is that the studied events are downstream of cellular entry. Manipulation will, necessarily, also interfere with invasion efficiency. To overcome this, we used conditions that still enabled pathogen entry and had a measurable effect on cocoon formation (see Methods). Furthermore, we focused only on *Shigella* that successfully invaded the host cytosol (galectin-3-positive BCV) as indicator for efficient infection. The readout rupture time can indicate changes in actin cocoon dynamics, but is not sufficient since it includes successive actin-regulated steps from cellular uptake to vacuolar rupture (Figure S2A). For this reason, we analyzed in parallel the percentage of bacteria per cell with actin cocoon (Figure 3B). We observed the strongest impact by the inhibition of the Arp2/3 complex with a 2.5-fold reduction of actin cocoons per cell (+CK666: 29.6±2.76%, p<0.0001). While Arp2/3 complex inhibition did not impede bacterial uptake, it was sufficient to prevent F-actin nucleation at the BCV (Figure 3B). Thus, the Arp2/3 complex regulates cocoon assembly. Interestingly, inhibition of PI3-kinases (42.3±0.41%, p<0.0001) or myosin II (38.1±6.92%, p<0.0001) significantly reduced the amount of actin cocoons (Figure 3B), and the remaining ones had decreased thickness and a less dense F-actin meshwork, respectively. On the other hand, inhibition of formins, ROCK, and the Rho GTPase Rac1 did not prevent cocoon formation (Figure 3B). Yet *Shigella* entered delayed into formin-inhibited cells (19.6±13.1 min, p=0.0007, Figure 3C) and we observed clearly reduced actin accumulation around the BCV. Eventually, formins might be rather involved in cocoon maintenance by F-actin elongation than filament nucleation. Cytochalasin D inhibits actin polymerization and was used as negative control for actin cocoon assembly. We barely detected actin rearrangements at the entry site and no cocoon-like structure after inhibition of actin polymerization (Figure 3B), underlining again that cocoon assembly requires *de novo* F-actin polymerization. We also observed a strongly reduced infection rate, probably caused by impaired actin rearrangements at the entry site and in line with the increased vacuolar rupture time (+DMSO: 16.4±11.4 min, +CytD: 39.6±19.3 min (p<0.0001)). Taken together, these results emphasize first a role of Cdc42-N-WASP-Arp2/3-mediated actin rearrangements around the vacuolar niche of *Shigella*. Second, preventing or perturbing actin cocoon formation at *Shigella*’s BCV impacts cytosolic access, underlining the importance of the cocoon for intracellular niche formation.

**Figure 3.**
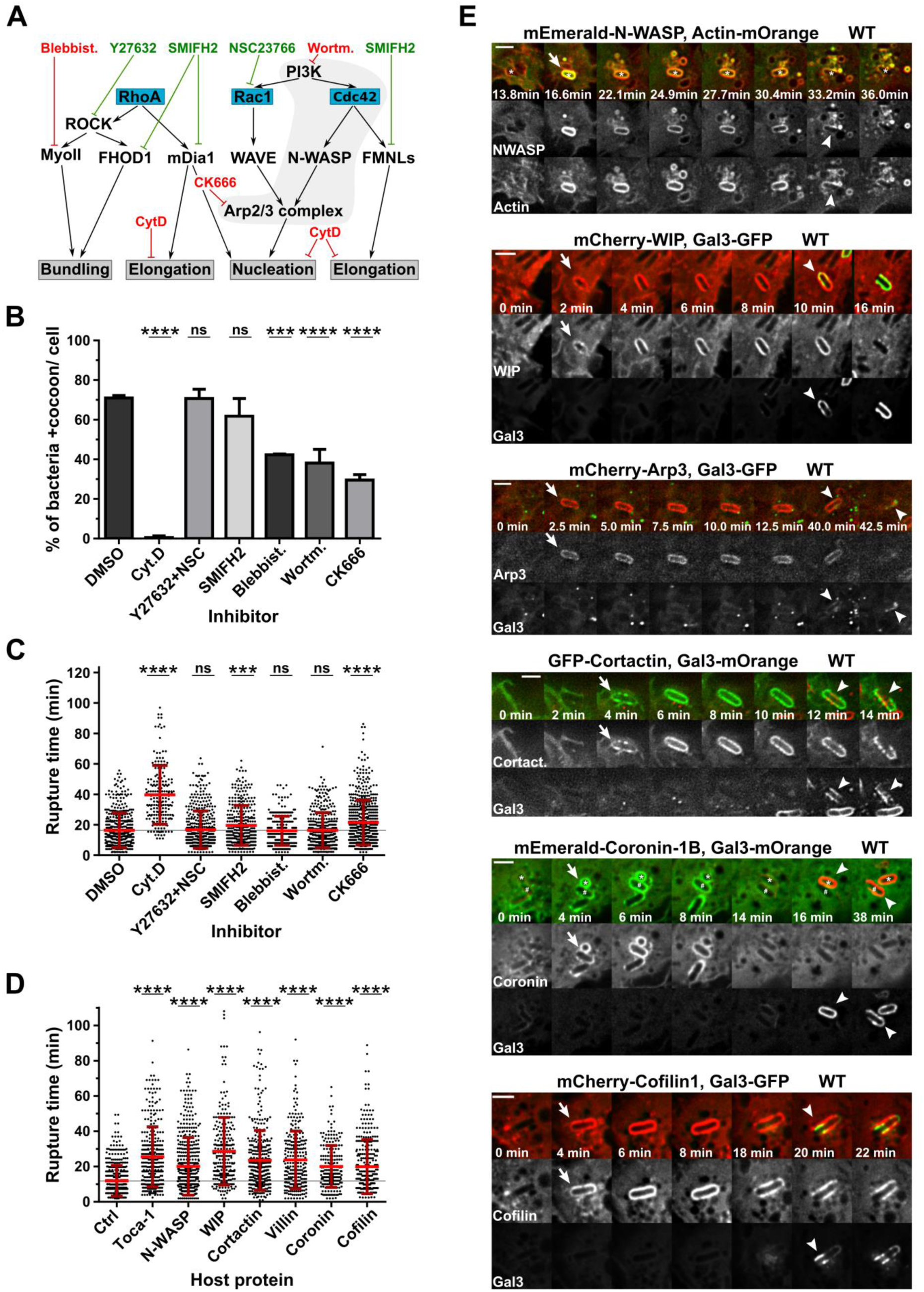
Host actin regulators are involved in actin cocoon assembly and recruited to *Shigella*’s BCV of before vacuolar escape. (**A**) Scheme of host proteins inhibited by compounds of the inhibitor screen (B-C). Blue box: Rho GTPases, black: ABPs and their regulators, grey box: effect on F-actin, red: inhibits cocoon assembly, green: no effect on initial cocoon assembly. CytD: Cytochalasin D, actin polymerization inhibitor; SMIFH2: formin inhibitor; NSC23766 and Y27632: Rac1 GEF and ROCK kinase inhibitors; Blebbist: Blebbistatin, myosin II ATPase inhibitor; Wortm: Wortmannin, PI3Ks inhibitor; CK666: Arp2/3 inhibitor. (**B-C**) Quantitative analysis of inhibitor screens identifies host proteins and signaling pathways involved in cocoon assembly (B) and regulation of cytosolic access (C) (n=3, N=2912 total invaders, on average 416 per condition; DMSO: Control). (**D**) Overexpression of seven selected host ABPs that interfere with the timing of vacuolar rupture (Ctrl: control, n≥3, N=2641, on average 330 per condition). (**E**) Recruitment of host ABPs to *Shigella*’s BCV before cytosolic escape. Time-lapses of HeLa cells expressing fluorescently-tagged host proteins and actin or galectin-3 (scale bars: 3 μm). All ABPs are recruited to the BCV (arrow) before vacuolar rupture (arrow head). Indicated are mean values ± SD, Student’s *t*-test was used with p<0.05 as significant compared to control (ctrl) (***p<0.001, ****p<0.0001, ns: not significant).

To identify host factors involved in actin cocoon formation, we overexpressed selected ABPs and their regulators. We hypothesized that their cellular excess interferes with F-actin polymerization and, in case these proteins are involved in cocoon regulation, potentially alters *Shigella*’s cytosolic access. We first screened ABPs for their recruitment to infection sites, determined afterwards the rupture time of positive candidates, and finally monitored their precise subcellular localization (Figure 3D,E). Remarkably, cytosolic escape of *Shigella* invading cells overexpressing Toca-1 (also known as FNBP1L), N-WASP, WIP1, Cortactin, Villin, Coronin-1B, or Cofilin-1 was significantly delayed (Figure 3D). N-WASP is an NPF for the Arp2/3 complex stabilized by WIP and activated by Cdc42-GTP, Toca-1, as well as PI(4,5)P_2_. Toca-1 activity itself is regulated by Cdc42 and PI(4,5)P_2_ (Rottner et al., 2017) (Ho et al., 2004) (Rohatgi et al., 1999). Cortactin binds to F-actin and the Arp2/3 complex and links signaling, cytoskeleton, as well as trafficking proteins to actin filaments (Kirkbride et al., 2011). Overexpression of these ABPs probably caused increased F-actin nucleation at the BCV membrane. Villin bundles F-actin, and its overexpression likely stabilizes actin cocoons. Further, coronin-1B collaborates with cofilin-1 in actin disassembly (Cai et al., 2008) (Chan et al., 2011). Overexpression of actin, coronin-1B, or cofilin-1 may boost F-actin turnover and polymerization at *Shigella*’s BCV. Since cocoons exhibited clear borders with a uniformly distributed, exceptional high fluorescence intensity (Figure 1A-C), and due to the identified ABPs (Figure 3), we expect a rather stiff, highly bundled and cross-linked F-actin meshwork.

Next, we performed time lapse microscopy of *Shigella* infections in cells co-transfected with the involved host proteins and galectin-3. We followed their recruitment to the BCV and correlated their localization with vacuolar rupture (Figure 3E). N-WASP was recruited to the *Shigella*-containing vacuole and co-localized there with actin. We did not observe any case of N-WASP recruitment to the BCV without cocoon assembly, or vice versa. In addition, WIP, the Arp2/3 complex component Arp3, and cortactin were recruited to the BCV membrane and all were, like actin (Figure 1C), at least partially depleted before vacuolar rupture (Figure 3E). Coronin-1B was recruited early during infection and dissociated from the BCV before appearance of the galectin-3 signal, while cofilin-1 localized around the vacuolar membrane remnant even after rupture (Figure 3E). Taken together, we identified a network of host actin regulators that are recruited to *Shigella*’s BCV and regulate cytosolic escape.

### Actin cocoon formation depends on Cdc42-activated and Arp2/3 complex-mediated F-actin nucleation

To verify the involvement of Cdc42 and the Arp2/3 complex in cocoon assembly, we performed siRNA-mediated knockdown experiments. Knockdown resulted in clearly altered entry sites with reduced membrane ruffling. Although Cdc42 knockdown or overexpression of its inactive mutant strongly reduces infection efficiency up to 75% (Mounier et al., 1999) (Mellouk et al., 2014), it does not entirely prevent cellular uptake in our experiments (Figure 4A,B, Figure S4B,C, S5A-E). Hence, we only considered bacteria that successfully invaded host cells to focus on the role of the actin cocoon. Strikingly, cocoon formation was completely or strongly abolished by knockdown of Cdc42 and ArpC3, respectively (+siRNA CDC42: 2.60±1.84%, +siRNA ArpC3: 16.6±0.47%, Figure 4B). We confirmed the strong delay of *Shigella* vacuolar rupture in Cdc42 and ArpC3 depleted cells (Figure S5D,E). Consequently, cocoon assembly depends on Cdc42-activated and Arp2/3 complex-mediated F-actin nucleation with Cdc42 signaling via N-WASP (see above). Thus, the main NPF of the Arp2/3 complex at the BCV is N-WASP. Interestingly, although knockdown of ARPC3 was very efficient, cocoon formation was not as strongly inhibited as for Cdc42 (Figure 4B). Either the remaining amount of Arp2/3 complex was sufficient to assemble some actin cocoons, or another, Cdc42-dependent actin polymerization pathway might be involved, for example via formins.

**Figure 4.**
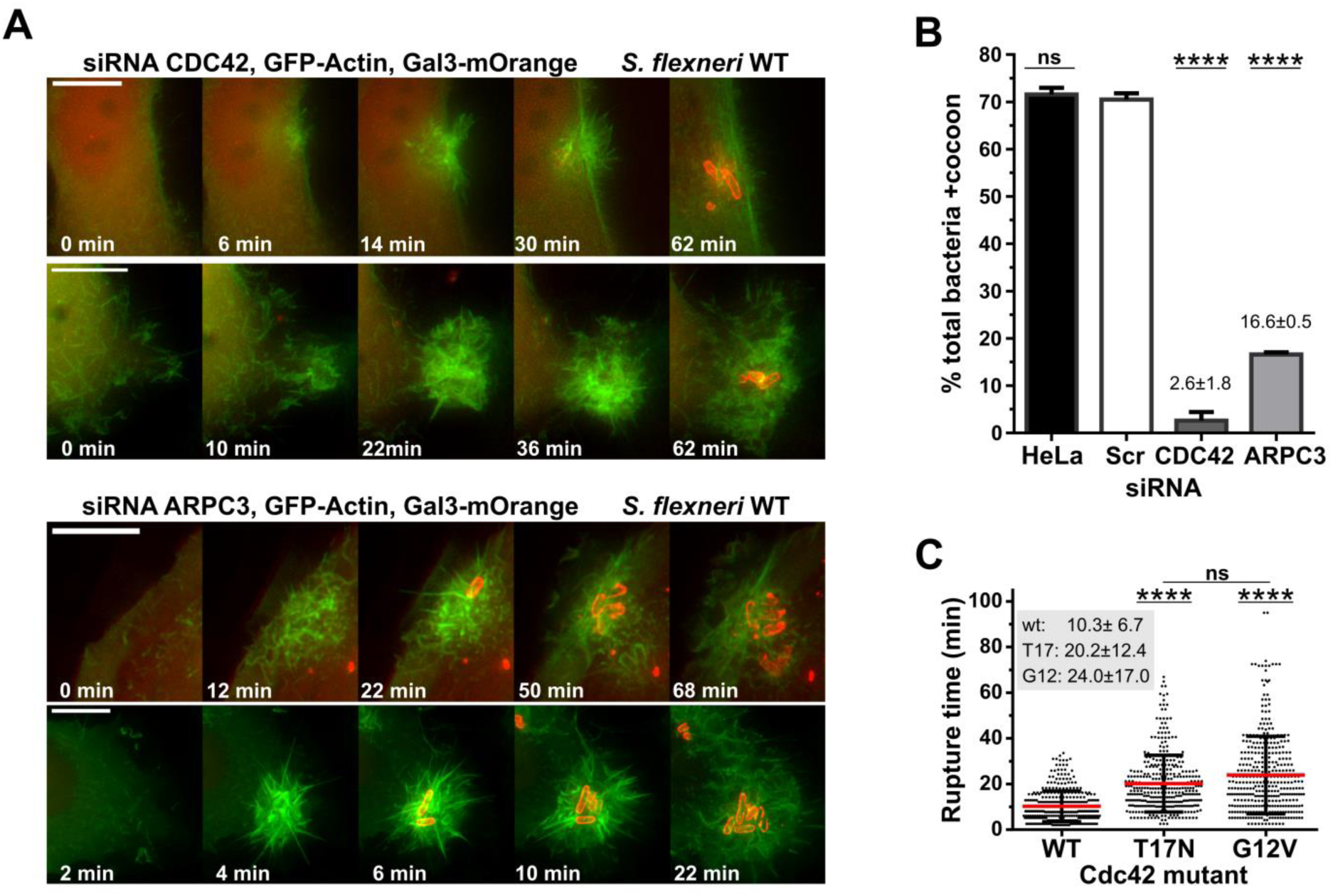
Cdc42-activated F-actin nucleation by the Arp2/3 complex is essential for cocoon formation. Knockdown of either CDC42 or the Arp2/3 complex subunit ARPC3 inhibits cocoon assembly (A-B). (**A**) Live microscopy monitoring actin rearrangements during early *Shigella* WT infection. Two representative time-lapses for CDC42 (top, KD 87.1%) and ARPC3 knockdown (bottom, KD 95.6%) are presented (scales: 5 μm, 1.05 μm z-stack, see Figure S4). Altered actin cytoskeleton rearrangements at the entry site indicate successful knockdown. (**B**) Quantitative analysis of actin cocoon assembly shows strongly reduced cocoon formation. Depicted are mean values ± SD (each n=3; no siRNA (HeLa): N=374, scramble siRNA (Scr): N=481, siRNA CDC42: N=464, siRNA ARPC3: N=573). (**C**) The ability of Cdc42 to cycle between an active and inactive state is required for efficient cytosolic access of *Shigella*. T17N: inactive, GDP-bound mutant, G12V: active, GTP-bound mutant (n=3, N=1123 total rupture events with WT: 373, T17N: 375, G12V: 375). Statistical significance: p<0.05 is significant compared to Scr (B) or WT (C) (****p<0.0001).

In addition, we investigated the dependence of Cdc42 activity on vacuolar escape by comparing the vacuolar rupture time of invading *Shigella* in cells overexpressing Cdc42 WT, an GDP- or GTP-bound Cdc42 mutant. Locking Cdc42 either in an active or inactive state significantly delayed the cytosolic escape of *Shigella* (Figure 4C, Figure S5I). Thus, cytosolic release of *Shigella* strongly depends on the ability of Cdc42 to function as molecular switch.

### Actin cocoon assembly is regulated by *Shigella* T3SS effector proteins

To identify which *Shigella* effectors are involved in actin cocoon regulation, we screened a *Shigella* mutant library with single T3SS effector deletions (Sidik et al. 2014). Results of nine selected mutants are presented (Figure 5). Since the amount of entering bacteria per cell varies for different mutants, we focused our analysis on the 1^st^ invading bacteria of individual strains. Strikingly, single deletions of the bacterial effectors IpgD, IpgB1 and IcsB strongly increased cytoskeletal rearrangements around the BCV. We found up to threefold more Δ*icsB* residing in an actin-containing structure before cytosolic escape (Figure 5A, Figure S6). We could also confirm our previous reported finding that IpgD increases cocoon formation of the total invading population (Mellouk et al., 2014). The rupture timing of Δ*ipgB1*, Δ*virA* as well as Δ*icsB* was likewise significantly delayed (Figure 5B). IcsA recruits N-WASP to the pole of cytosolic bacteria for actin tail formation (Suzuki et al., 1998) (Egile et al., 1999), but is not accessible before vacuolar rupture. Consistent with this, we observed no effect on cocoon assembly in Δ*icsA* infections (Fig.5A). Thus, IpgD, IpgB1, and IcsB play a role in actin cocoon regulation. Deletion of these *Shigella* effectors might lead to a misregulation of the cocoon with a disturbed actin turnover.

**Figure 5.**
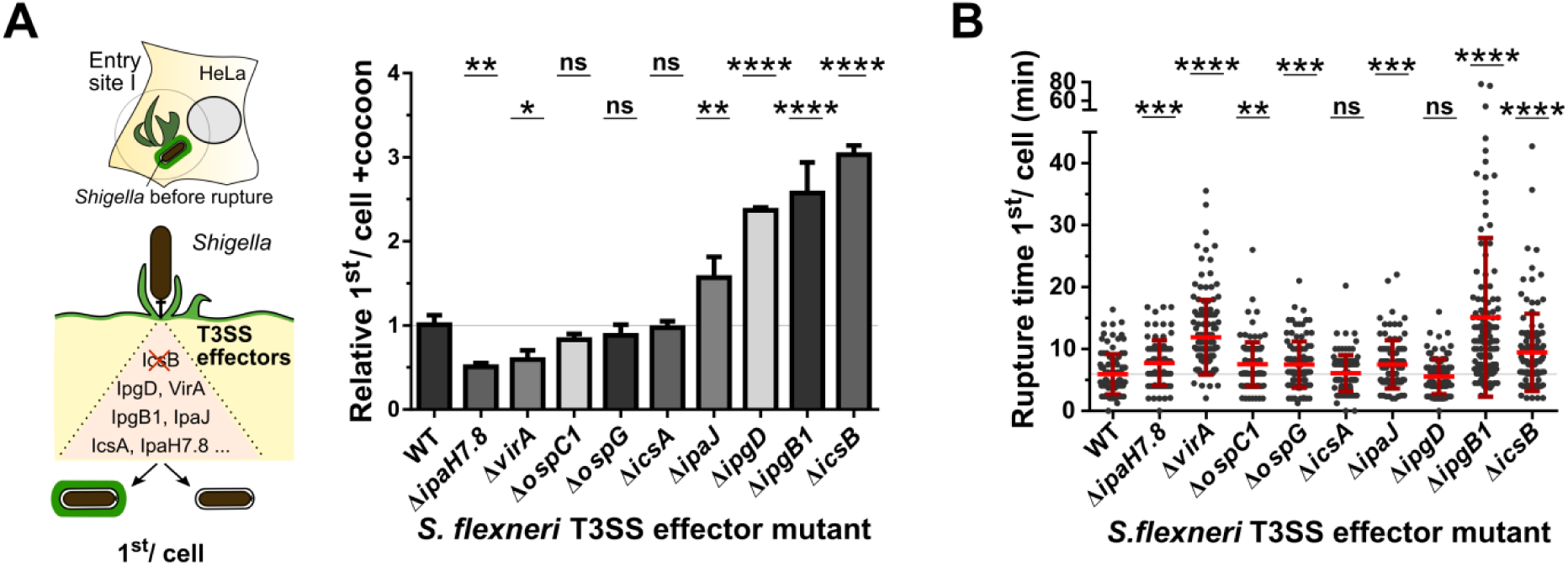
The actin cocoon and cytosolic access are regulated by *Shigella* T3SS effector proteins. HeLa cells expressing actin-GFP and galectin-3-mOrange were infected with *Shigella* single effector deletion mutants. Several deletions had strong effects indicating involvement in the regulation of actin cocoon formation (**A**) and vacuolar escape (**B**). N=1085 first invading bacteria per cell, WT: N=101, Δ*ipaH7.8*: N=80, Δ*virA*: N=131, Δ*ospC1*: N=112, Δ*icsA*: N=103, Δ*ipaJ*: N=95, Δ*ipgD*: N=121, Δ*ipgB1*: N=116, Δ*icsB*: N=120. Indicated are mean values ± SD of n≥3 independent experiments normalized to *Shigella* WT. p<0.05 is significant compared to WT (*p<0.05, **p<0.01, ***p<0.001, ****p<0.0001, ns: not significant). (See Figure S4, S5)

### Clustering of host actin regulators at *Shigella*’s BCV depends on the T3SS effector IcsB, but not their initial recruitment

We finally questioned if the recruitment of the actin nucleation machinery at the BCV (Figure 3E) depends on specific *Shigella* effectors. We focused on IcsB, the strongest hit of our T3SS effector screen (Figure 5). During canonical phagocytosis, active Cdc42 localizes to forming pseudopods, but gets inactivated and depleted to complete cup closure (Niedergang and Grinstein, 2018) (Freeman and Grinstein, 2014) (Hoppe and Swanson, 2004). Likewise, Cdc42 initiates *de novo* F-actin actin rearrangements for *Shigella*’s cellular uptake (Adam et al., 1996). We confirmed Cdc42 recruitment to forming membrane ruffles during *Shigella* infection. Yet contrary to canonical phagocytosis, Cdc42 remained at the BCV after cellular uptake and persisted with galectin-3 at the membrane remnant (Figure 6A,B). Remarkably, we deciphered that this constant localization of Cdc42 depends on IcsB (Figure 6C,D). Deletion of IcsB did not interfere with initial Cdc42 recruitment to membrane ruffles, but lead to its depletion from the BCV before vacuolar rupture (Figure 6C). While in WT *Shigella* infections 91.9±1.56% of cytosolic bacteria constantly localized Cdc42 at the BCV, deletion of IcsB dramatically impaired Cdc42 clustering by 13-fold (Figure 6D). As for Cdc42, Toca-1 also clustered IcsB-dependent at *Shigella*’s vacuole following initial recruitment (Figure 6E). Noteworthy, while the membrane remnant was quickly recycled in WT *Shigella* infections, it remained loosely and sticky around cytosolic Δ*icsB* bacteria (Figure 6C,E).

**Figure 6.**
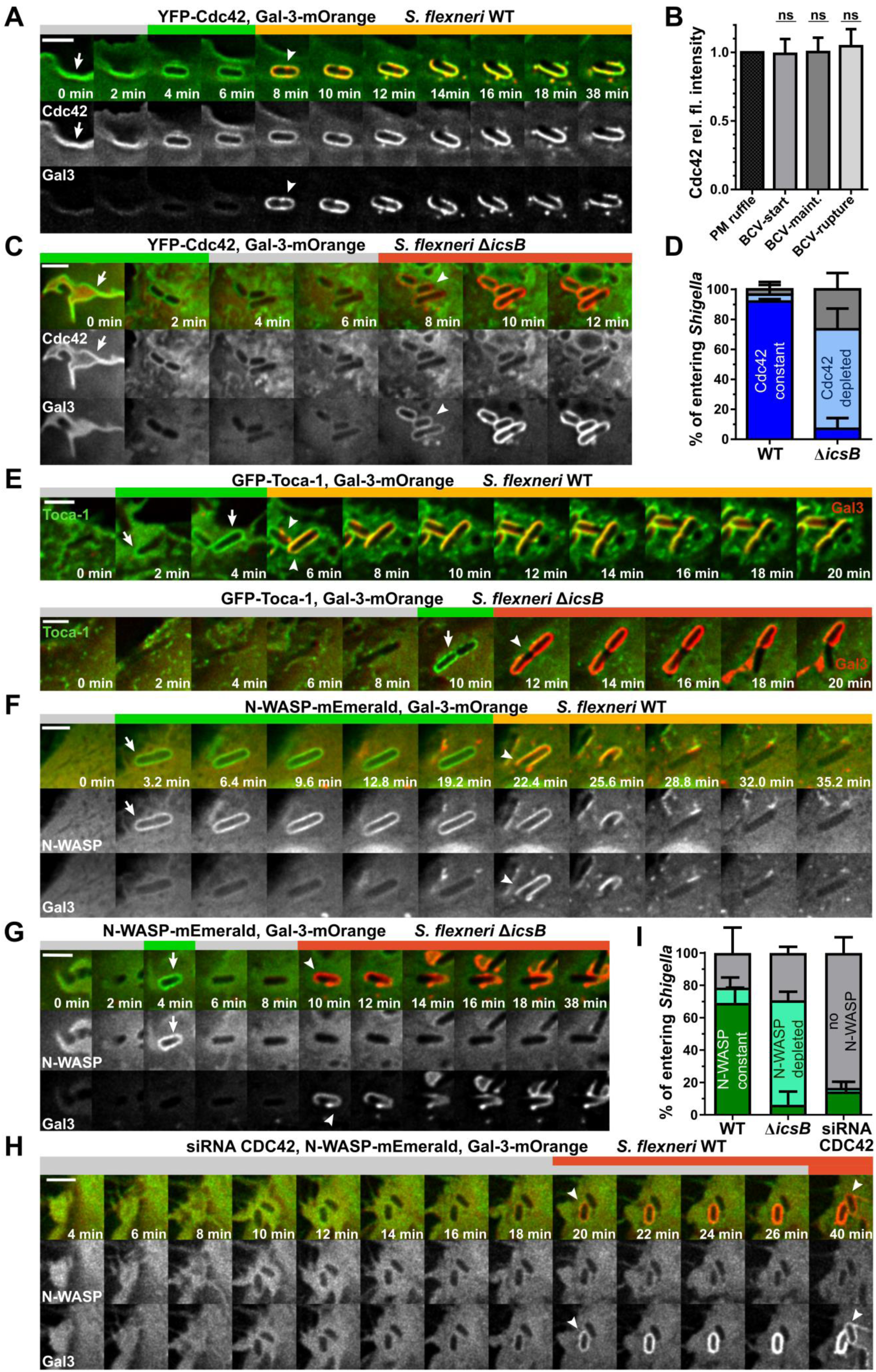
The constant localization of host actin regulators at the BCV depends on the *Shigella* effector IcsB, but Cdc42 is crucial for initial N-WASP recruitment. (**A**) Representative time-lapses of constant Cdc42 localization at the BCV during *Shigella* WT infections. (**B**) Quantification of (A) with normalized relative fluorescence intensity of Cdc42. PM ruffle: Fluorescence intensity at plasma membrane ruffles, BCV-start: at the phagocytic cup before scission, BCV-maint: at maintained BCV after cellular uptake, BCV-rupture: at BCV after rupture. (**C**) Deletion of IcsB leads to Cdc42 removal from the BCV before rupture, but does not prevent its initial recruitment to membrane ruffles. (**D**) Quantitative analysis of successfully invading *Shigella* with either constant (dark blue), transient (light blue), or no (grey) Cdc42 localization at the BCV (time-lapses (A,C), n=3, N=311). *Shigella* WT vs. Δ*icsB*: constant Cdc42 p<0.0001, depleted Cdc42 p<0.001, no Cdc42 not significant (ns). (**E**) Toca- 1 is likewise recruited to and constantly localized at the BCV during *Shigella* WT infection (top), while it gets depleted before rupture in Δ*icsB* infections (bottom). (**F-G**) The *Shigella* effector IcsB constantly localizes N-WASP at the BCV during infection. One representative time-lapse for *Shigella* WT (F) and Δ*icsB* (G) are shown. Initial N-WASP recruitment is IcsB- independent (B). (**H**) Cdc42 knockdown strongly impairs initial N-WASP recruitment to the BCV of *Shigella* WT. (**I**) Quantification of N-WASP localization at the BCV of invading *Shigella* with either constant (dark green), transient (light green), or no (grey) N-WASP (time-lapses 7A-C, n=3, N=441). Statistical significance: *Shigella* WT vs. Δ*icsB*: constant N-WASP localization p<0.0001, N-WASP depleted p<0.0001, no N-WASP ns. *Shigella* WT vs. siRNA Cdc42: constant N-WASP localization p<0.0001, N-WASP depleted ns, no N-WASP p<0.0001. Statistical significance: one-way ANOVA (B) or two-way ANOVA with Tukey’s multiple comparisons test (D,I). Arrow: initial protein recruitment, arrow head: moment of vacuolar rupture, scale bars: 3 µm. Green bar: protein at BCV, red bar: Galectin-3 at BCV, yellow bar: protein and galectin-3 at BCV remnant, grey bar: no protein or galectin-3.

Finally, we found N-WASP constantly localized around *Shigella*’s endocytic compartment in Caco-2 (Figure S7B) and HeLa cells. N-WASP was recruited after cellular entry to 78.6±3.25% of WT bacteria that successfully escaped from the BCV, and remained associated with the BCV membrane remnants in 69.2±9.45% of the cases (Figure 6F,I). As for Cdc42, the permanent localization at the BCV, but not the initial recruitment of N-WASP was IcsB-dependent (Figure 6G). We further confirmed N-WASP recruitment to be independent of the *Shigella* virulence factor IcsA (Figure S7A). We hypothesized that Cdc42 initiates N-WASP recruitment, as Cdc42 localizes during very early entry steps at membrane ruffles. Therefore, we knocked down Cdc42 and followed the localization of N-WASP during *Shigella* WT infection (Figure 6H). Interestingly, we found a strong reduction in initial N-WASP recruitment to the BCV (16.67±7.92%), but recruited N-WASP remained localized (14.48±4.84%, Figure 6H,I). We concluded that residual Cdc42 activity was sufficient to recruit N-WASP in some rare cases with constant BCV localization due to IcsB. Thus, we found that IcsB clusters key factors of the host actin nucleation machinery at the BCV. But the initial recruitment of the Arp2/3 complex activator N-WASP depends on Cdc42 signaling. Since knockdown of Cdc42 prevents cocoon formation, the Cdc42-mediated regulation of N-WASP is crucial for actin cocoon assembly.

## DISCUSSION

We identified in this study highly dynamic *de novo* F-actin polymerization of exceptional thickness around the endocytic vacuole of intracellular *Shigella*. We showed that this actin cocoon assembles after scission at the surface of the entire vacuolar membrane, and does not constitute remaining F-actin from cell entry. Compared to previous reported actin rearrangements around internalized phagosomes, we found that the cocoon shares features with the “actin flashing” phenomenon that occurs target- and cell-unspecific around endocytic compartments to protect cells from overloading degradative pathways (Yam and Theriot, 2004) (Liebl and Griffiths, 2009). Actin flashing has been defined as fast, successive waves of transient actin polymerization and depolymerization at membranes of fully internalized, early phagosomes (Yam and Theriot, 2004) (Liebl and Griffiths, 2009). Both phagosomal actin rearrangements in common is (i) the involvement of certain host proteins like Arp2/3 (Bierne et al., 2001) (Yam and Theriot, 2004), (ii) assembly at the cell periphery and (iii) preferably at later formed phagosomes. Besides, *Shigella*’s actin cocoon is clearly distinct and resembles a longer-lasting structure with non-cyclical actin turnover that is much denser than any other cellular actin structure. Its disassembly leads to immediate cytosolic escape of *Shigella*, while actin flashing is followed by vesicle fusion for phagosome maturation (Liebl and Griffiths, 2009). Therefore, assembly of these distinct actin structures precedes completely different phagosomal fates. We propose that *Shigella* hijacks the host actin flashing mechanism for its own needs and thus counteracts a cellular response to invasion.

Despite this, the endocytic vacuole of *Shigella* is intrinsically different from canonical phagosomes with regard to its molecular composition. We identified a network of host actin regulators that are recruited and prolonged localized at the BCV (Figure 3, 7). Interestingly, *Shigella* plays itself an important role in actin cocoon regulation by inducing localized signaling at with its effectors IpgD, IpgB1, and IcsB. One strategy to locally induce actin rearrangements is to immobilize key proteins by either (i) increasing their membrane binding affinity, or (ii) preventing their removal. Both strategies imply subverted host protein localization by *Shigella*. We found IcsB to be sufficient to cluster Cdc42, Toca-1 and N- WASP at *Shigella*’s vacuole (Figure 7). Strikingly, it has recently been discovered that IcsB posttranslationally modifies these membrane-bound actin regulators by lipidation (Liu et al., 2018). Yet the biological function of this modification remained unknown. Our findings place IcsB as central *Shigella* effector for the re-localization and entrapment of signaling proteins to regulate cytosolic access. IcsB most likely lipidates its substrates directly at the BCV membrane, since it localizes around intracellular bacteria (Baxt and Goldberg, 2014) (Campbell-Valois et al., 2015) (Liu et al., 2018) (Figure 7B). Thus, *Shigella* seems to “glue” host proteins for localized signaling to its endocytic vacuole by adding an additional membrane binding motif. The precise mechanistic consequences of this modification on cocoon formation remain to be further investigated.

**Figure 7.**
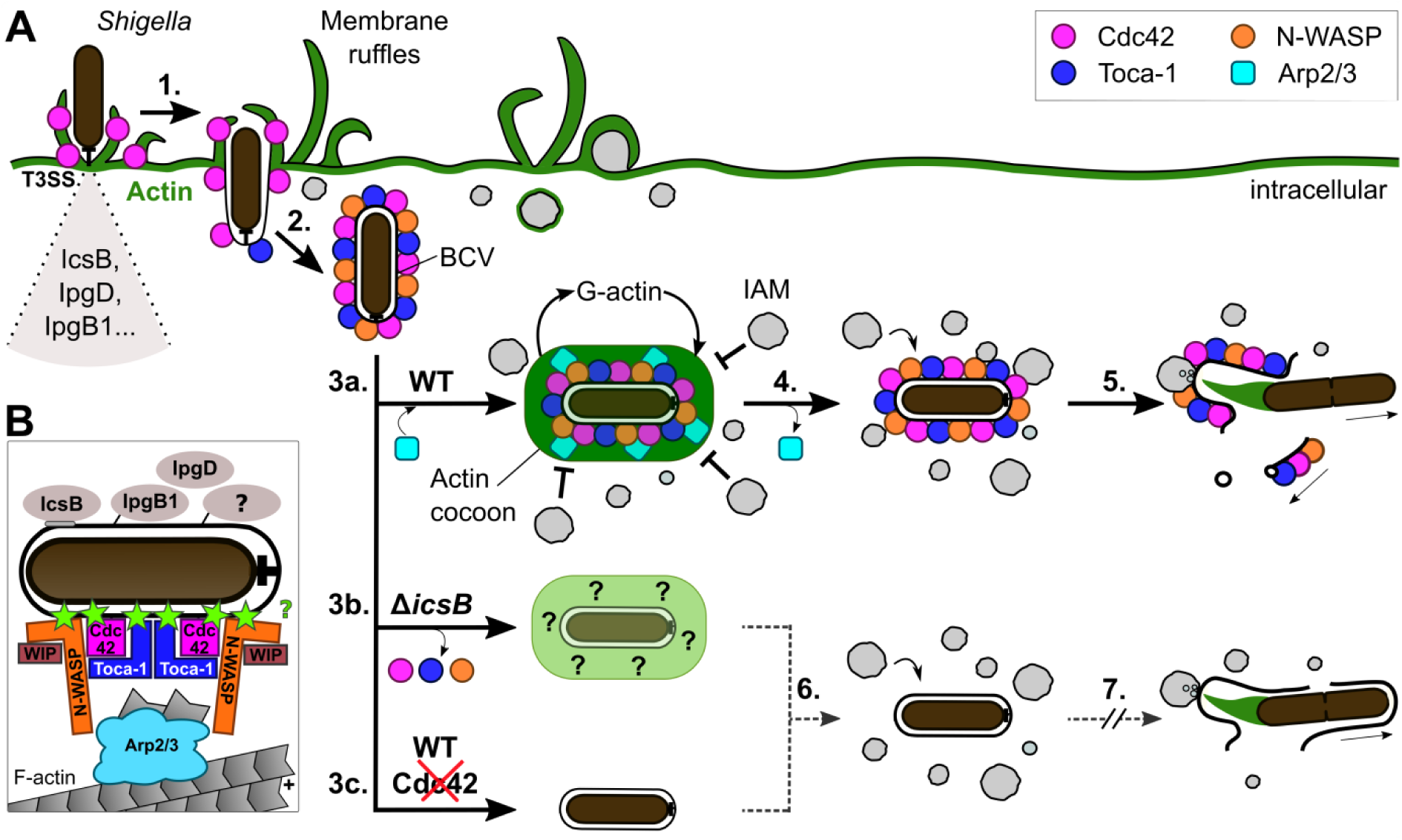
Model for *Shigella*-mediated subversion of host actin cytoskeleton to regulate niche integrity. (**A**) Injected *Shigella* T3SS effectors subvert a network of host proteins for actin cocoon regulation. First, Cdc42 is recruited during membrane ruffling at the forming BCV (1). After scission, Toca-1 and N-WASP localize around the entire BCV and cluster there with Cdc42 (2). The Arp2/3 complex is recruited and the actin cocoon assembles *de novo* (3a). The cocoon is maintained with constant G-actin turnover and shields the BCV from surrounding vesicles like IAMs. During cocoon disassembly, the Arp2/3 complex gets depleted from the BCV, while Cdc42, N-WASP and Toca-1 remain at the BCV (4). Cocoon disassembly is followed by vacuolar rupture and *Shigella* escape into the host cytosol for replication, while Cdc42, N-WASP and Toca-1 remain at the recycled BCV membrane remnant (5). In the absence of IcsB, Cdc42, N-WASP and Toca-1 are not immobilized at the BCV, leading to an altered, actin-containing cytoskeletal structure (3b). Depletion of Cdc42 impairs initial N-WASP recruitment and thus actin cocoon formation (3c). Both, disturbing cocoon regulation (3b) or preventing its formation (3c) interferes with and delays vacuolar rupture (6, 7). (**B**) Components of the immobilized host actin nucleation machinery and identified *Shigella* effectors that regulate cocoon formation. Green star: constant, IcsB-dependent localization at the BCV (Figure 6, 7) and reported (Liu et al., 2018) posttranslational lipidation.

Interestingly, IcsB is not required for initial N-WASP localization at the BCV, but Cdc42 (Figure 6, 7). This reveals an unexpected second function of active Cdc42 during infection besides cellular uptake: It is required for efficient actin cocoon assembly, which in turn regulates cytosolic access. However, Cdc42 recruitment and activation probably involves other effectors targeting upstream Rho signaling during cellular entry. For instance, IpaC activates Cdc42 via Src signaling (Tran Van Nhieu et al., 1999, Mounier et al., 2009, Adam et al., 1996), IpgB1 is a membrane-bound Rho GEF at intracellular *Shigella* (Ohya et al., 2005, Huang et al., 2009, Weigele et al., 2017), and the inositol-phosphatase IpgD produces PI5P from PI(4,5)P_2_ (Niebuhr et al., 2002) (Pendaries et al., 2006). Consistent with this we identified both, IpgB1 and IpgD, as actin cocoon regulators (Figure 5).

Furthermore, we deciphered that actin cocoon dynamics need to be tightly regulated. Cocoon disassembly precedes cytosolic escape, which places it as gatekeeper for successful invasion. Perturbing proper actin cocoon regulation by interfering with either cocoon assembly or disassembly consequently diminishes cytosolic invasion of *Shigella*. Remarkably, cocoons around the Δ*ipgB1* and Δ*icsB* mutants were more pronounced, highlighting the interference of these effectors with actin turnover. Impairing the activation and immobilization of Rho GTPases by deletion of these effectors probably causes a misregulation of the actin cocoon ultrastructure with disturbed actin dynamics. We found that the Δ*icsB* BCV differs in its host protein composition from WT *Shigella*. We propose that this induces the formation of a changed cytoskeletal structure that is less prone to cytosolic escape (Figure 7). As *Shigella* subverts the host actin cytoskeleton via distinct pathways, deletion of major regulators puts an imbalance in this complex, fine-tuned system.

It emerges that several human pathogens assemble varying actin structures around their intracellular niches. It is tempting to speculate that, as in cells, different structures result in different functions. New actin rearrangements have been associated with maintaining vacuolar stability, trafficking to acquire nutrients, or preventing phagosome maturation. Often, components of additional cytoskeletal systems and F-actin assemble during late infection steps around the replicative niche of pathogens with long vacuolar lifestyles (Kumar and Valdivia, 2008) (Meresse et al., 2001) (Sukumaran et al., 2011). For example, the *Salmonella*-induced, loose actin meshwork around vacuolar bacteria stabilizes the phagosome and maintains vacuolar integrity (Meresse et al., 2001) (Vazquez-Torres et al., 2000) (Poh et al., 2008) (Unsworth et al., 2004). Contrary, the actin cocoon is clearly distinct, indicating a different function. Yet how this new actin structure fulfills its function on a precise mechanistic level is unclear.

We found that cocoon-surrounded BCVs were stationary, excluding a role in actin-driven motility of endocytic compartments (Merrifield et al., 1999) (Taunton et al., 2000). Second, a function in sorting and recycling as indicated by dot-like actin structures (Puthenveedu et al., 2010) (Seaman et al., 2013) seems unlikely, since actin assembled homogenously around entire BCVs. Third, we never detected fusion events of dextran-positive vesicles with the BCV, nor recruitment of markers for phagosome maturation (Mellouk et al., 2014) (Weiner et al., 2016) (Kühn et al., 2017), ruling out a function in vesicle fusion. We previously demonstrated that the actin cocoon forms between the BCV and surrounding vesicles (Weiner et al., 2016). Therefore, we suggest that the cocoon acts as “gatekeeper” to regulate pathogen entry into the host cytosol. It shields the BCV as physical barrier from the host endomembrane system to prevent fusion with lysosomes. Interestingly, Cdc42-N-WASP-Arp2/3 have been reported to prevent phagosome maturation by an F-actin coat around the replicative niche of *Leishmania donovani* (Lodge and Descoteaux, 2005). Additionally, the actin cocoon could prevent host recognition and non-canonical LC3- associated phagocytosis, because a role of IcsB in autophagy escape by preventing LC3 recruitment to the BCV vacuole has been reported (Campbell-Valois et al., 2015) (Ogawa et al., 2005) (Baxt and Goldberg, 2014). As cocoon inhibition also delays vacuolar escape, cocoon formation itself allows proper vacuolar maturation required for the rupture mechanism. The cocoon and its involved molecular machinery could support either vacuolar rupture or its membrane recycling by governing short distance endomembrane trafficking.

In conclusion, we discovered a novel microbial subversion strategy: The activation and immobilization of the host actin nucleation machinery by posttranslational modification causes localized signaling at the *Shigella* vacuole during early infection. This leads to the formation of an exceptional, thick actin structure to ensure niche integrity and cytosolic escape (Figure 7). The dynamic maturation of this structure has to be tightly regulated as either preventing or disturbing it impairs cytosolic escape.

## ACKNOWLEDGMENTS

For tools and fruitful discussions we are grateful to John Rohde, Emmanuel Lemichez, Amel Mettouchi, Christian Merrifield, Guy Tran Van Nhieu, Sandrine Etienne-Manneville, Claude Parsot, Serge Mostowy, Yuen-Yan Chang and Patricia Latour-Lambert. We thank Noelia López-Montero and Laurent Audry for technical support, and members of the BCI department for critical discussions. This work was supported by grants from the ERC (EndoSubvert) to J.E., the ANR to J.E. (StopBugEntry) and to C.Z. (ANR-16-CE16–0019-01, NEUROTUNN). J.E. is a member of the LabEx consortium IBEID and MilieuInterieur. S.K. is a recipient of a Pasteur-Roux fellowship.

## AUTHOR CONTRIBUTIONS

Conceptualization, Project Administration: S.K., J.E.; Methodology, Visualization: S.K.; Investigation: S.K., J.B.; Formal Analysis: S.K., J.B.; FRAP analysis: S.B; Resources: J.E., C.Z.; Writing – Original Draft: S.K.; Writing – Review & Editing: S.K., J.E.; Supervision: S.K., L.B., J.E.; Funding Acquisition: S.K., C.Z., J.E. All authors discussed the results and commented on the manuscript.

## METHODS

### Bacterial strains

The following streptomycin-resistant *Shigella flexneri* 5a M90T-Sm (GenBank #CM001474.1) derived strains harboring the pWR100 virulence plasmid (GenBank #AL391753.1) were used: wild type M90T (WT), M90T (WT) expressing the protein afimbrial adhesin Afa-I (afaE1 gene) from *E. coli* (Labigne-Roussel et al., 1984), M90T (WT) expressing dsRed and AfaI (Yuen-Yan Chang), and M90T Δ*icsB*-2 (FXS299) (Claude Parsot). The screened library of *S. flexneri* single deletion T3SS effector mutants was kindly provided by John R. Rohde (Dalhousie University) as part of a pwR100 collection originating from M90T (WT) AfaI (Sidik et al., 2014). We used *Salmonella enterica* serovar Typhimurium (*S.* Typhimurium) strain SL1344 pGG2-dsRed (WT) expressing dsRed. The *E. coli* InvA strain containing the *Yersinia* invasin A (Isberg et al., 1987) and *S. flexneri* M90T (WT) expressing mCherry and Afa-I (*E. coli* plasmid pIL22) were provided by Guy Tran Van Nhieu. *S. flexneri* strains were grown in trypticase soy broth (TCSB) with 50 μg/ml ampicillin, and cultured at 37 °C on TCSB agar including 0.01% congo red to select for functional T3SS system. *S.* Typhimurium was grown at 37 °C in lysogeny broth (LB) medium supplemented with 0.3 M NaCl and 50 μg/ml ampicillin, while *E. coli* InvA was cultured in LB medium with 100 μg/ml ampicillin.

### Cell culturing, bacterial infection and immunocytochemistry

Human epithelial HeLa cells (clone CCL-2, ATCC), HeLa cells stably expressing galectin-3- mOrange (Patricia Latour-Lambert), and intestinal epithelial Caco-2 TC7 cells (kindly provided by P. Sansonsetti) were cultured in Dulbecco’s modified Eagle’s medium (DMEM, Thermo Fisher Scientific) supplemented with 10% (v/v) heat-inactivated fetal bovine serum (FBS) at 37 °C, 5% (HeLa) or 10% (Caco-2) CO_2_. Cell lines were checked negative for mycoplasma. HeLa cells were used for all experiments except Figure S1B and S7B. All infection assays were performed in EM buffer (120 mM NaCl, 7 mM KCl, 1.8 mM CaCl_2_, 0.8 mM MgCl_2_, 5 mM glucose, 25 mM HEPES, pH 7.3). Actin cocoon formation and vacuolar escape was not affected by EM buffer compared to FluoroBrite™ DMEM (Thermo Fisher Scientific) supplemented with 10% heat-inactivated FBS and 4 mM GlutaMAX.

For *Shigella*, infection cultures in TCBS plus antibiotics were inoculated with a 1:100 dilution of an overnight culture and incubated at 37 °C to an optical density (OD 600 nm) of 0.6-0.7. Then, bacteria were washed in PBS and resuspended in EM buffer. Strains without AfaI were incubated for 15 min with poly-L-lysine (Sigma) as described previously (Weiner et al., 2016). In general, bacterial dilutions were prepared for the following MOIs: live imaging 96-well format MOI 5-150 (normally MOI in range 15-40 to exclude effects of phagocytic load), live imaging 35 mm glass bottom µ-Dishes (#81158, Ibidi) MOI 15, poly-L-lysine treated bacteria MOI 50. Other bacterial pathogens, like *Salmonella* or *E. coli* InvA, were grown similarly in their corresponding medium. *Salmonella* infections at MOI 50 were started maximal 10 min after spinning down the bacteria in pre-heated Eppendorf tubes and washing in pre-heated EM buffer. *E. coli* InvA were added to imaged cells at an MOI of 50. For fixed experiments, *Shigella* (WT) or dsRed (WT) adhered for 10 minutes at 20 °C (MOI 50), before incubation at 37 °C for 45 min or 60 min. Samples were washed three times with PBS, fixed in cold 2% PFA, and stained with Alexa Fluor™ 647 Phalloidin (#A22287, Thermo Fisher) for 45 min.

### Plasmids, inhibitors, siRNAs, and transfection

The following plasmids were kindly provided by our colleagues: Arp3-pmCherryC1, Cortactin- pmCherryC1, as well as Cofilin-pmCherryC1 from Christian Merrifield (Taylor et al., 2011), Cdc42-mCherry, Cdc42-V12-GFP, as well as Cdc42-N17-GFP from Sandrine Etienne-Manneville, pEYFP-C1-Villin from Sylvie Robine (Revenu et al., 2007), pEGFP-C1-Toca-1 from Jennifer Gallop, pGFP-Cortactin from Kenneth Yamada (Addgene plasmid #50728), mEmerald-N-Wasp-C-18 and mEmerald-Coronin1B from Michael Davidson (Addgene plasmids #54199, 54050), pmCherry-C1-WIP from Anna Huttenlocher, (Addgene plasmid #29573), and pEYFP-Cdc42 from Joel Swanson (Hoppe and Swanson, 2004); Addgene plasmid #11392). The plasmids pEGFP-C3-actin, pOrange-C3-actin, pEGFP-N1-Galectin-3, and pOrange-Galectin3 have been described previously (Mounier et al., 2009) (Ehsani et al., 2012) (Mellouk et al., 2014). All plasmid constructs were verified by sequencing. For quantitative screening experiments (Figure 1F-G, 3B-D, 4C, 5; Figure S2C-D, S3, S5F-I), HeLa cells were seeded at a density of 7000 cells/well 24 h prior to transfection into 96-well plates (Greiner, #655090). Cells were then transfected with the respective plasmids using X- tremeGENE™ HP DNA transfection reagent (Roche, #6365779001) for 48 h (24 h for Toca-1 and Cdc42).

In inhibitor screens, the following compounds and concentrations (DMSO stock solutions dissolved in EM buffer) were applied: 1 µM Cytochalasin D (Enzo Life Sciences Inc, #BML-T109-0001), 10 µM SMIFH2 (Sigma-Aldrich, #S4826), 15 µM Y-27632 (BD biosciences, #562822), 50 µM NSC23766 (Calbiochem, #553502), 15 µM (±)-Blebbistatin (BioVision, #BV-2405-5), 10 µM Wortmannin (Enzo Life Sciences Inc, #BML-ST415-0001), 200 µM CK666 (Sigma, #SML0006). For each inhibitor, working concentrations were identified that did not affect cell viability in the duration of the experiment, but showed clear effects on either cell shape or actin rearrangements during infection. In general, cells were pre-incubated for 40 min at 37 °C in EM buffer containing the corresponding inhibitor. Afterwards, bacteria were added in EM buffer with the same inhibitor concentration and imaged immediately. For Cytochalasin D, cells were not pre-incubated and the inhibitor was added with the bacteria.

Following siRNAs were used in siRNA transfections: 50 nM SEPT7 (ID #s2743, Ambion), 10 nM ON-TARGETplus SMARTpool siRNAs from Dharmacon for CDC42 (#L- 005057-00-0005), ARPC3 (#L-005284-00-0005), as well as Non-targeting pool as negative control (#D-001810-10-05). For quantitative analysis (Figure 2D, 4B; Figure S5A-E), HeLa cells were seeded at a density of 5000 cells/well 24 h prior to transfection into 96-well plates. Reverse siRNA transfection was performed using the Lipofectamine RNAiMAX Transfection Reagent (Invitrogen, #13778030) for 72 h. After 48 h, transfection mixes were removed and cells were transfected for 24 h with pEGFP-C3-actin and pOrange-Galectin3 plasmids using X-tremeGENE™ HP DNA transfection reagent. In parallel, upscaled samples for Western blot quantification were prepared in 6-well plates and for live imaging at the DeltaVision wide- field microscope in 35 mm glass bottom µ-Dishes (Ibidi, #81158) (Figure 4A, 6H,I).

### Light microscopy

For quantitative screening experiments (Figure 1F-G, 2D, 3B-D, 4B-C, 5; Figure S2C,D, S3, S5), image acquisition was performed using an inverted epifluorescence Nikon Ti-E wide field microscope with a 20x (0.5 Numerical Aperture (NA), 2.1 Working Distance (WD)) N- Plan air objective, an automatic programmable xy stage, and the Nikon perfect focus system. Cells were imaged every 1-2 min for 2-3 h inside a 37 °C heating chamber. *Shigella* (WT) infections were monitored as control in each experiment. To detect host protein recruitment to *Shigella*-induced endocytic compartments (BCV, IAMs), time-lapse movies of high resolution were recorded at 37 °C on a DeltaVision Elite (GE Healthcare) widefield microscope using a 60×/1.42 NA oil objective and a step size of 0.25-0.35 μm (Figure 1A-C, 2A-C, 3E, 4A, 6; Figure S1C, S6, S7). Imaging was performed for 2-3 h after adding the bacteria and the frequency of image acquisition was adjusted to the dynamics of the investigated event. Images were subsequently deconvolved using an integrated deconvolution analysis software (DeltaVision Elite). Bleach correction of illustrated images was performed using Fiji (http://fiji.sc). Fixed samples were imaged using a Perkin Elmer UltraView spinning disc confocal microscope, with a 60X×/1.3 N.A. oil objective and a Z step size of 0.3 μm (Figure S1A,B).

### FRAP measurement and data analysis

Live-cell FRAP experiments (Figure 1D,E) were performed at an inverted Perkin Elmer UltraView VOX confocal spinning disk microscope equipped with a FRAP module and Volocity software. Images were acquired using a 60X×/1.3 N.A. oil objective with a single Z plane. Cells expressing actin-GFP in 35 mm glass bottom culture dishes were imaged at 37 °C in EM buffer. Cells were infected with *Shigella* (WT) DsRed and bacterial presence was controlled before bleaching of each single cocoon structure. We monitored the fluorescence of GFP-actin using low intensity laser excitation (488 nm) (pre-bleach scans). A circular region was photobleached with the same laser excitation at high intensity (decrease of the fluorescence into the ROI by 60-80%), and fluorescence recovery monitored over time (post-bleach scans). 10 pre-bleach images were recorded at 1 s intervals, followed by a single bleaching pulse. One cocoon per cell was bleached with only half-bleached ones considered for analysis. The duration of post-bleach imaging was adjusted to the duration of fluorescence recovery. Stress fibers were bleached away from focal adhesions. Thus, actin turnover inside the filaments is expected to result from incorporation of unbleached actin from the cytoplasmic pool. Raw data were fitted with simFRAP plugin in Image J. For each acquisition reference cell, photobleached cell and bleached area are selected to extract the mobile fraction F_m_. Data were normalized to the first post-bleaching step and plotted and analyzed in Prism version 6 (GraphPad software).

### Antibodies and Western blotting

Primary antibodies used were polyclonal rabbit anti-SEPT7 (IBL, #18991, 1:2000), and monoclonal mouse anti-β-actin (Sigma-Aldrich, #A5316, 1:2000), anti-Cdc42 (BD Transduction Laboratories, #610929,1:500), as well as anti-ArpC3 p21-Arc (Clone26, #612234, BD Transduction Laboratories, 1:1000). Secondary antibodies were anti-mouse (Bio-Rad, #170-6516, dilution 1:1000 or 1:2000) and anti-rabbit (Bio-Rad, #170-6515 dilution 1:2000) horseradish peroxidase-conjugated. To quantify knockdown efficiency by immunoblotting, siRNA treated HeLa cells were lysed with RIPA buffer containing protease inhibitor (Roche, #11836170001) for 30 min at 4 °C. Protein quantification of lysates was done with the Micro BCA™ Protein Assay Kit (Thermo Fisher Scientific, #23235) and 10 µg of total protein was loaded into NuPAGE™ 4-12% Bis-Tris Protein Gels (Thermo Fisher Scientific, #NP0321BOX). Proteins were transferred to a nitrocellulose membrane using the Trans-Blot® Turbo™ RTA Mini Nitrocellulose Transfer Kit and Trans-Blot© Turbo™ Transfer System (Bio-Rad). Antibody detection was carried out with the SuperSignal™ West Pico PLUS Chemiluminescent Substrate (Thermo Fischer Scientific, #34577) with actin as loading control.

### Quantitative and statistical image analysis

Time lapse microscopy series were analyzed using Icy (http://icy.bioimageanalysis.org/), Volocity 6.3 (PerkinElmer), or Fiji (http://fiji.sc) to determine starting points of foci formation, actin cocoon assembly, actin cocoon lifetime, vacuolar rupture time, and localization of host proteins. At least three biologically independent experiments (n) per condition were performed. Quantitative data are mean values with error bars indicating ± Standard Deviation (SD) if not mentioned differently. Statistical analysis was performed in GraphPad Prism version 6. If not indicated differently, statistical significance was determined using a two- tailed Student’s *t*-test (Mann-Whitney) or a one-way ANOVA followed by Dunnett’s multiple comparison test. (ns) not significant, p<0.05 was considered as significant: *p<0.05, **p<0.01, ***p<0.001, ****p<0.0001. For additional information on data analysis see Supplementary Methods.

## SUPPLEMENTAL INFORMATION

Supplementary Information includes 7 figures and Supplementary Methods and can be found online with this article.

