## Supplemental Information for "A new form of actin assembly around the *Shigella*-containing vacuole regulates its intracellular niche"

#### Inventory of Supplemental Information

Figure S1, related to Figure 1
Figure S2, related to Figure 1
Figure S3, related to Figure 1
Figure S4, related to Figure 2, 4, and 7
Figure S5, related to Figure 4
Figure S6, related to Figure 6
Figure S7, related to Figure 6

Supplementary Experimental Procedures

SUPPLEMENTAL FIGURES

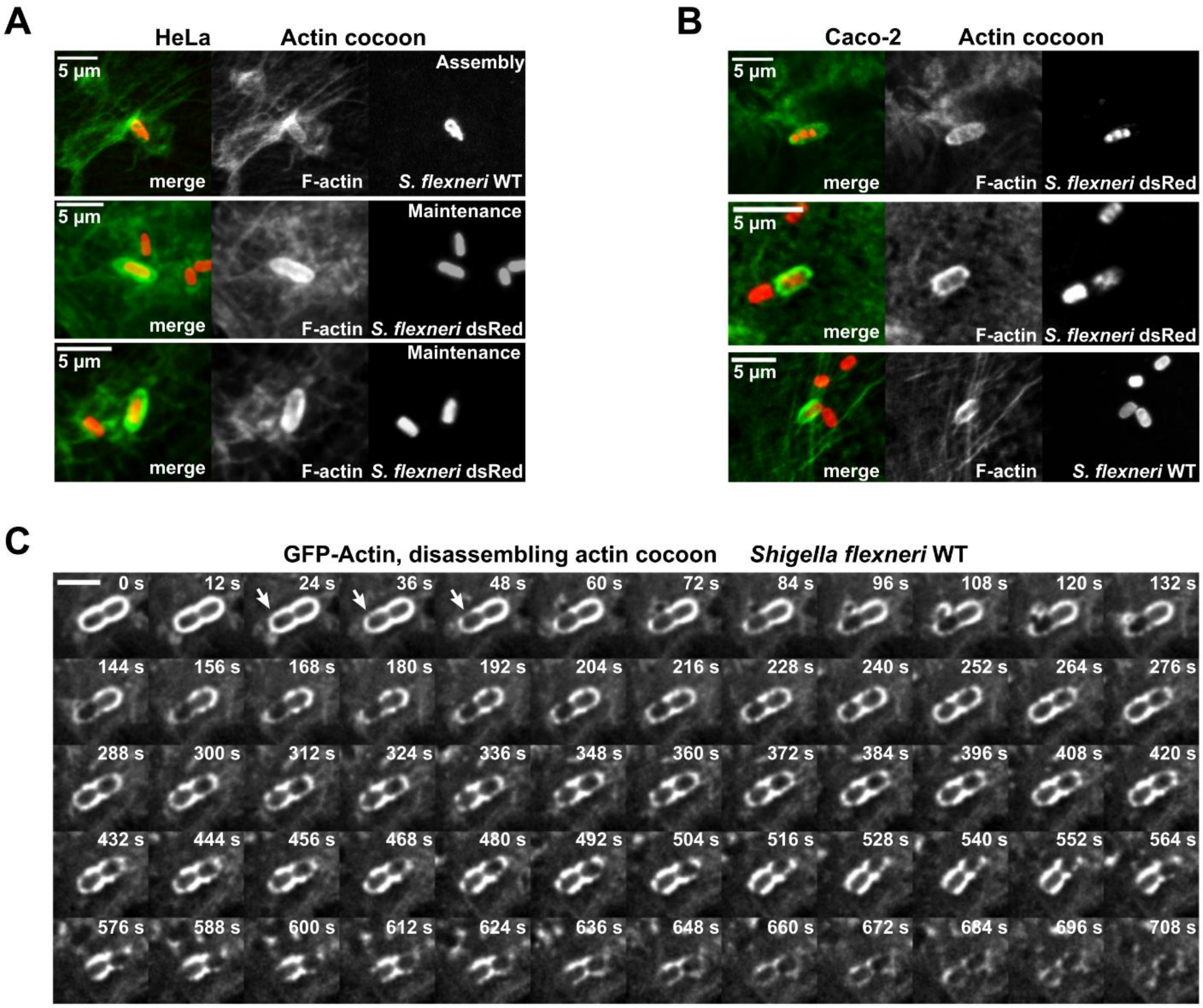


**Figure S1. Actin cocoons contain F-actin with constant turnover until final disassembly**
(**A,B**) Immunofluorescence images of HeLa (A) and Caco-2 cells (B) expressing endogenous actin. Cells were infected with DsRed-expressing *Shigella* WT strain or *Shigella* WT (red) and phalloidin-labeled for F-actin (green). (**C**) Real-time images showing that actin cocoons disassemble simultaneously at different locations while undergoing constant reassembly during *Shigella* WT infection in actin-GFP expressing HeLa cells (scale bar: 3 µm). (related to Figure 1)

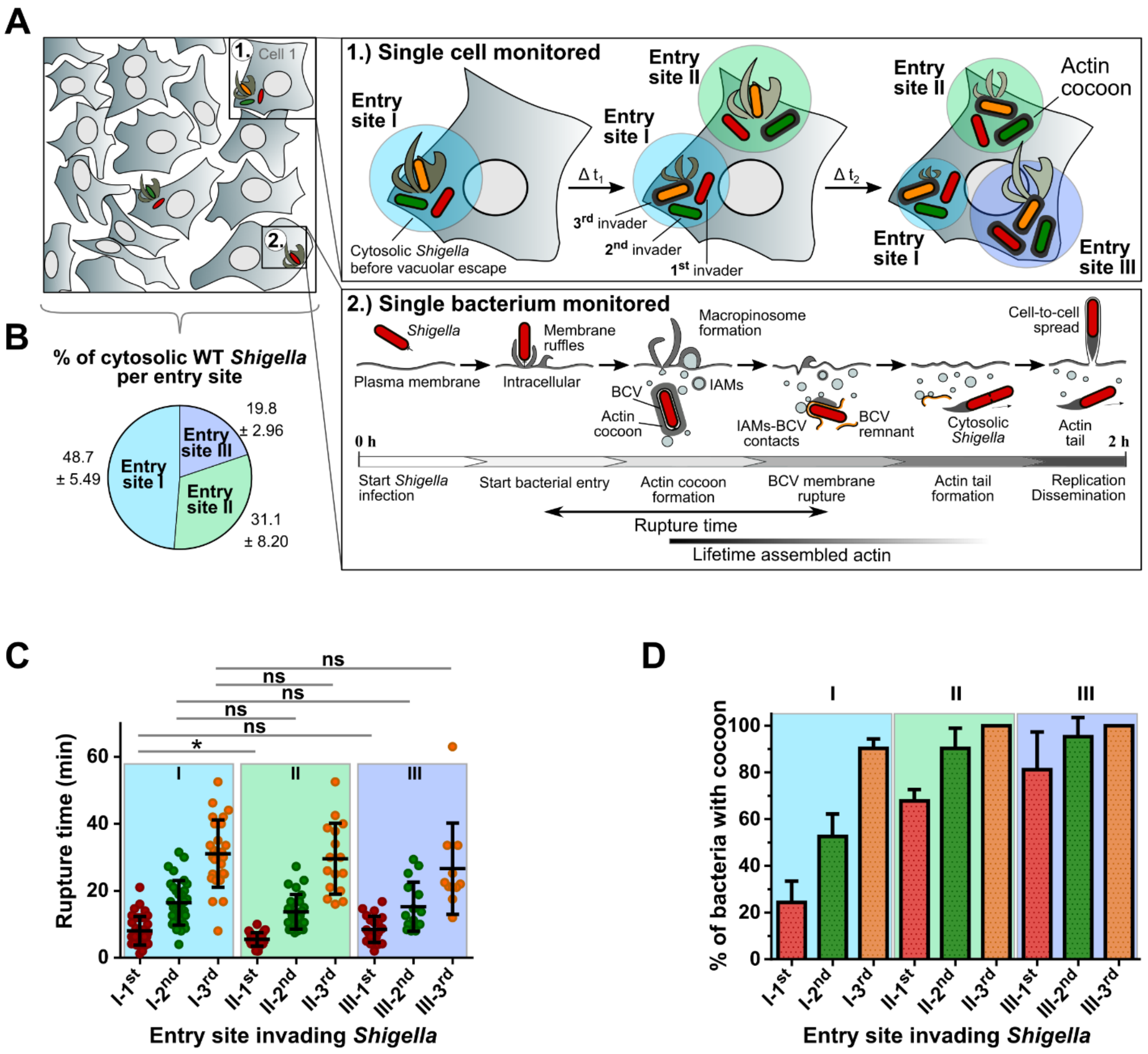


**Figure S2. Experimental setup to monitor successive invasion steps of *Shigella* in real time**
(**A**) Successive steps of invading *Shigella* are monitored simultaneously at single cell (1) and single bacterium (2) levels. This allows correlating them within the cellular context and with other invading bacteria. In this setup (at MOI of 40; Figure S3), single cells were infected on average via three (3.03±0.46) distinct entry sites per cell (I, II, and III). The 1$^{st}$ (red), 2$^{nd}$ (green), and 3$^{rd}$ (orange) invader of each entry site were considered in our analysis. (**B**) Almost half of the cytosolic *Shigella* WT entered host cells via the 1$^{st}$ entry site, while about 1/3 bacteria invaded via the 2$^{nd}$ and 1/5 via the 3$^{rd}$ foci. (**C,D**) Actin cocoon assembly and rupture time depend on entry site and order of infection. Depicted is the in-depth analysis of *Shigella* WT that developed an actin cocoon from figure 1F,G (n=4, N=446). The rupture time of individual 1$^{st}$, 2$^{nd}$, or 3$^{rd}$ bacteria invading the same cell through different entry sites is similar (C). The probability of actin cocoon formation for individual bacteria depends on the order of infection and the specific entry site (D). All late invading bacteria polymerize an actin cocoon. All 3$^{rd}$ invaders of foci II and III in all independent experiments had an actin cocoon. Error bars indicate ± SD (C), Mann-Whitney test with $p<0.05$ as significant (*$p<0.05$, ns: not significant). In (D), mean values ± SD are indicated. (related to Figure 1)

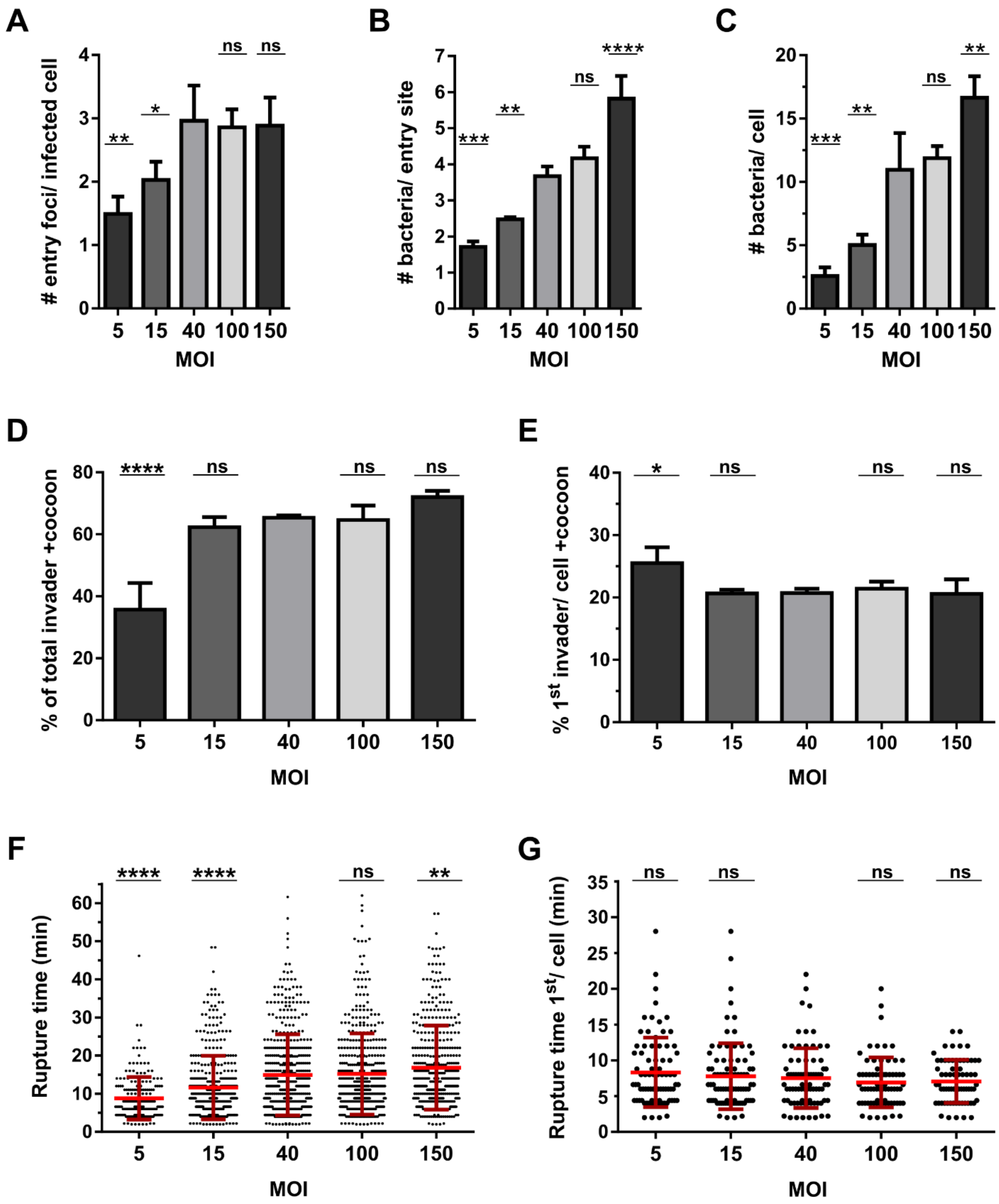


**Figure S3. The multiplicity of infection (MOI) defines the amount of entering *Shigella* per cell, but not the probability of actin cocoon formation**

The effect of MOIs from 5 to 150 was investigated with regards to entry sites per infected cell (**A**), successfully invading bacteria per entry site (**B**), overall number of invading bacteria per cell (**C**), the probability of cocoon formation (**D,E**), and vacuolar escape (**F,G**). The higher the MOI, the more bacteria enter via the same entry site (B). The quickly saturated effect of the MOI to cocoon formation (D) and rupture time (F) is probably due to the increased probability for cocoon assembly of later entering bacteria

(Figure S2). Since with high MOI more bacteria per entry site invade host cells, there are more late invaders present that assemble an actin cocoon. This probably increases the percentage of the total cytosolic *Shigella* with cocoons at higher MOIs compared to very low ones. Comparing the same bacterial population, here the 1st invading *Shigella* per cell, leads to no difference in actin cocoon formation and vacuolar escape (E,G). In (D,F), the total invading *Shigella* populations were analyzed (n=3, N=2000 bacteria with MOI 5: N=203, MOI 15: N=430, MOI 40: N=477, MOI 100: N=461, MOI 150: N=429)). In (E,G), the 1st invaders per cell of previous experiments were taken into account (N=385 with MOI 5: N=81, MOI 15: N=86, MOI 40: N=78, MOI 100: N=72, MOI 150: N=68). Indicated are mean values ± SD as error bars. Statistical significance was determined using one-way ANOVA or Student's *t*-test comparing individual MOI experiments with MOI 40. $p<0.05$ was considered as significant (*p<0.05, **p<0.01, ***p<0.001, ****p<0.0001, ns: not significant). (related to Figure 1)

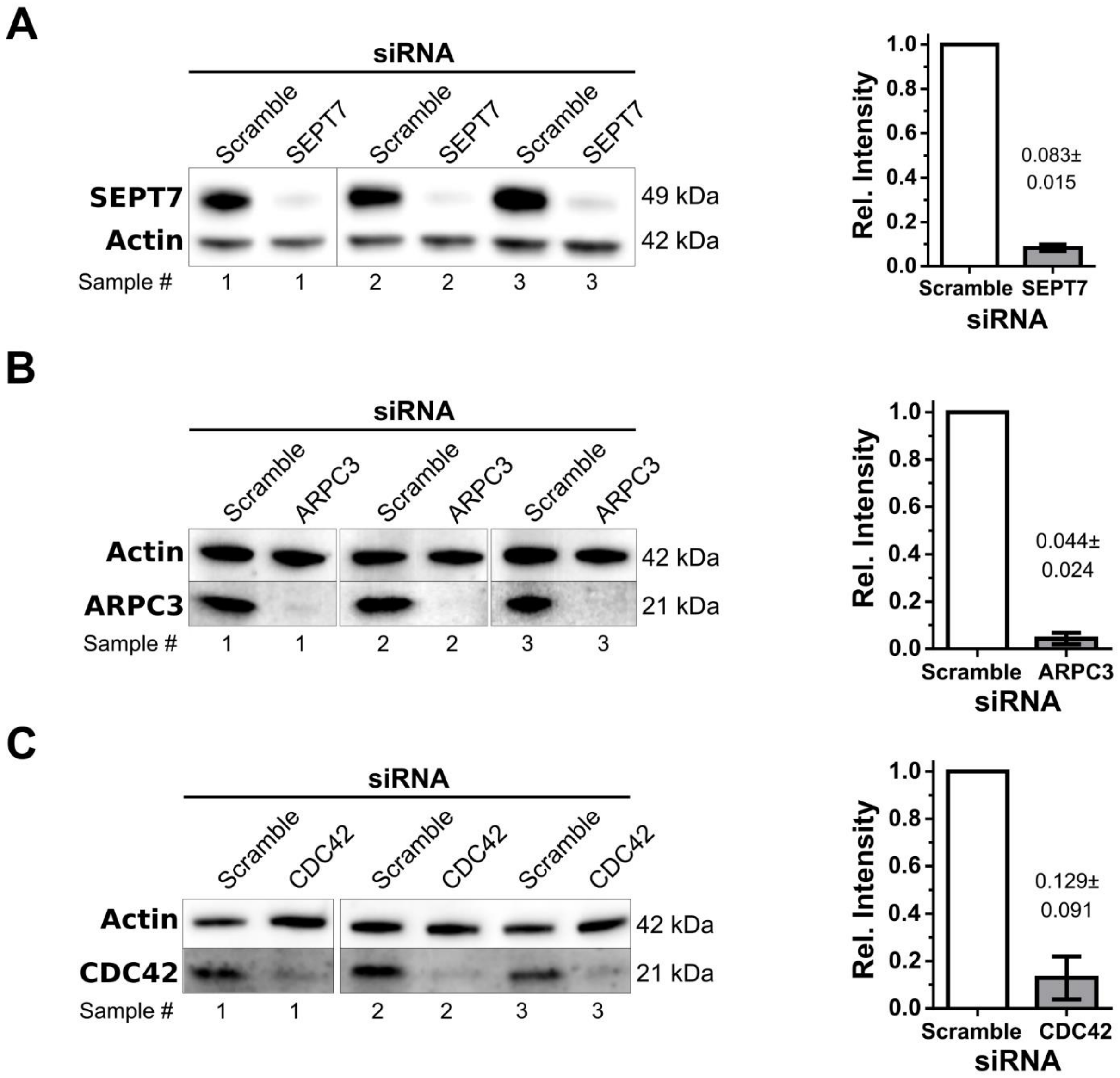


**Figure S4: Western Blot validation of knockdown experiments by RNA-interference.**
siRNA treatments against Septin-7 (SEPT7, **A**), the Arp2/3 complex component ARPC3 (**B**), and CDC42 (**C**). Samples of n=3 independent experiments are shown. Quantification of unmodified images revealed knockdown (KD) efficiencies of 91.7% (SEPT7, **A**), 95.6% (ARPC3, **B**), and 87.1% (CDC42, **C**). The contrast in the figures was enhanced to highlight KD efficiency. (related to Figure 2, 4, 7)

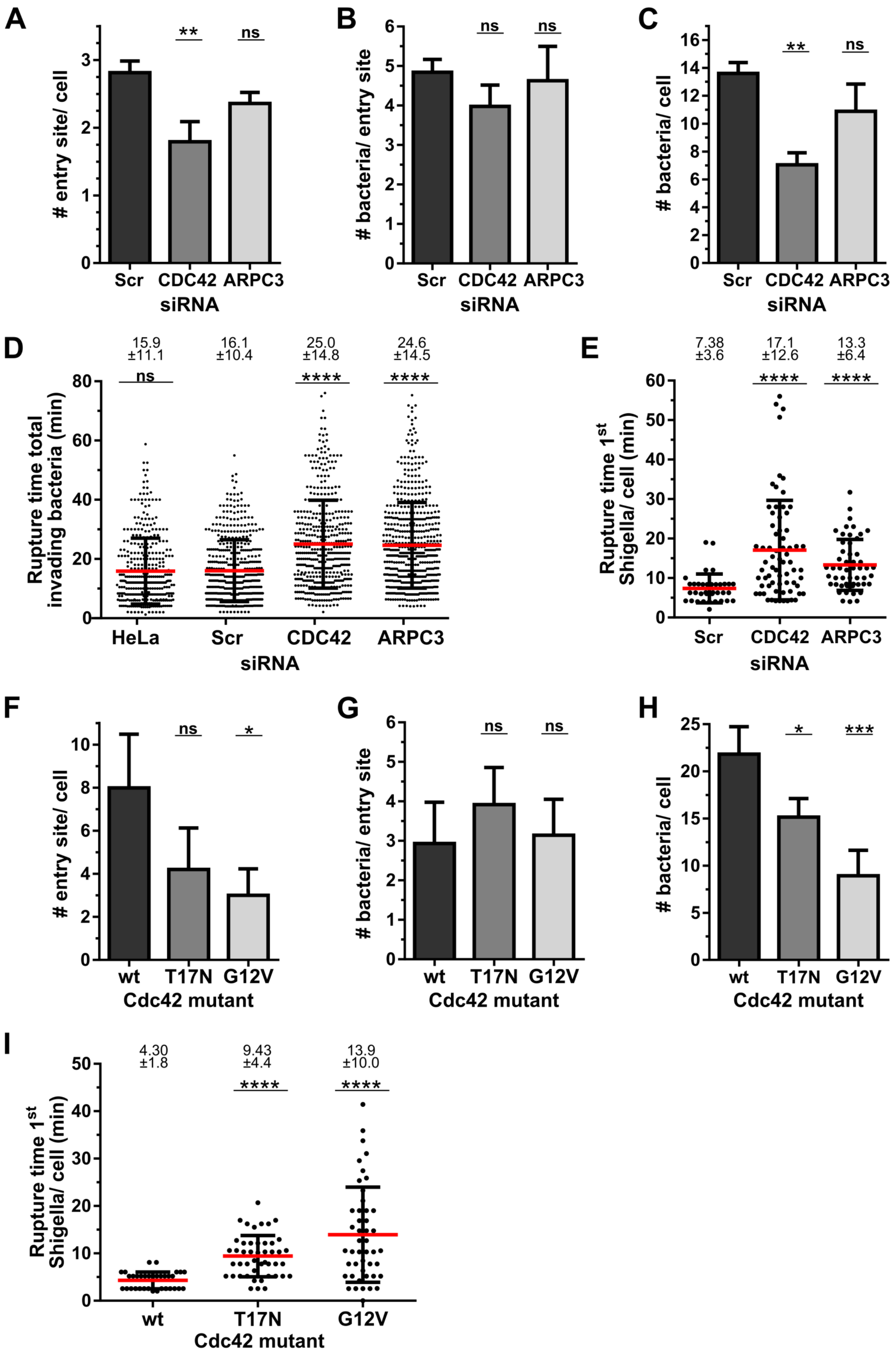
A
### entry site/ cell
**
ns
Scr
CDC42
ARPC3
siRNA
B
### bacteria/ entry site
ns
ns
C
### bacteria/ cell
D
15.9 ±11.1
16.1 ±10.4
25.0 ±14.8
24.6 ±14.5
****
Rupture time total invading bacteria (min)
HeLa
E
7.38 ±3.6
17.1 ±12.6
13.3 ±6.4
Rupture time 1st Shigella/ cell (min)
F
*
wt
T17N
G12V
Cdc42 mutant
G
H
***
I
4.30 ±1.8
9.43 ±4.4
13.9 ±10.0

**Figure S5: Cdc42 and the Arp2/3 complex are involved in host cell invasion and vacuolar rupture.** Depicted are additional statistics for the experiments shown in figure 4. Bacteria that successfully escaped into the host cytosol were considered for quantitative analysis. (**A-E**) Knockdown efficiency was confirmed by immunoblotting (Figure S4). Decreasing cellular levels of Cdc42 decreased the number of entry sites per infected cell (A) and thus the number of invading bacteria per cell (C). Depicted are the rupture times for all invading bacteria per experiment (D) or the first invaders per cell (E). (**F-I**) HeLa cells coexpressing Galectin-3 and Cdc42 WT, T17N or G12V were infected with *Shigella* WT. Presented are number of entry sites per cell (F), number of bacteria per entry site (G), total number of invading bacteria per cell (H), and rupture time of the first invading *Shigella* per cell (I). Depicted are error bars indicating ± SD, or mean values ± SD. One-way ANOVA or Student's *t*-test was used to determine significance compared to infected control or Cdc42 WT-expressing cells. $p<0.05$ was considered as significant (*$p<0.05$, **$p<0.01$, ***$p<0.001$, ****$p<0.0001$, ns: not significant). Counted events: (A-D) N=1892 with no siRNA (HeLa): N=374, control scramble siRNA (Scr): N=481, siRNA Cdc42: N=464, siRNA ArpC3: N=573; (e) N=157 with scramble: N=36, siRNA Cdc42: N=69, siRNA ArpC3: N=52; (F-H): N=1123 with WT: N=373, T17N: N=375, G12V: N=375; (I): N=128 with WT: N=34, T17N: N=47, G12V: N=47. (related to Figure 4)

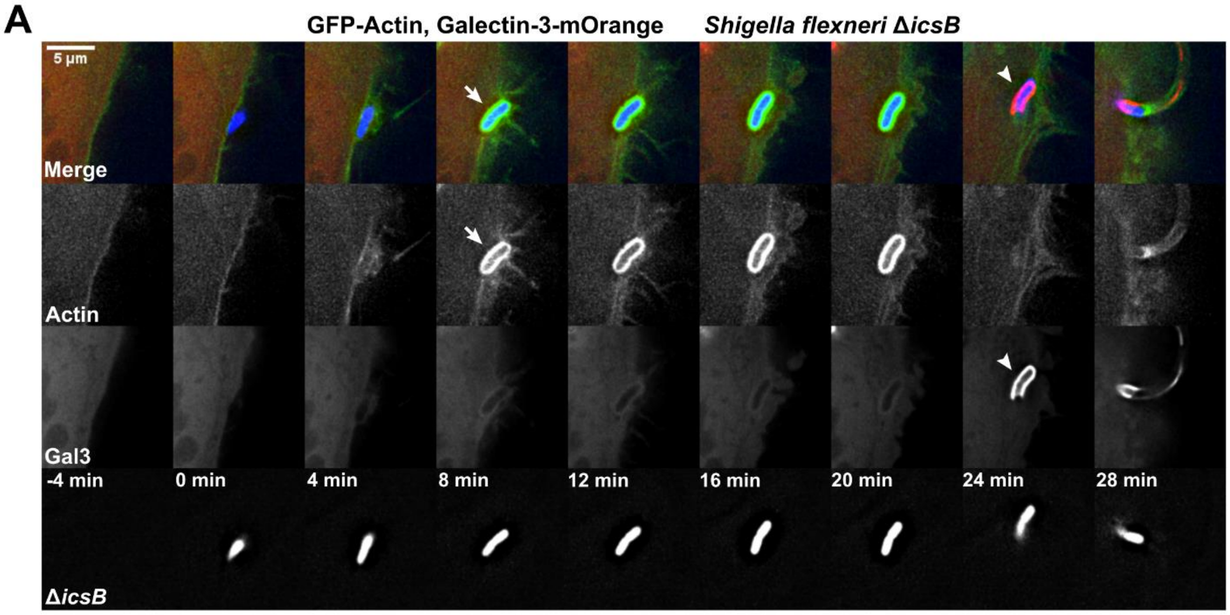


**Figure S6. Actin rearrangements at the BCV of *Shigella* Δ*icsB***

(**A**) Representative time-lapses of HeLa cells co-transfected with actin and galectin-3 and infected with *Shigella* Δ*icsB* DsRed. t=0 min: start entry site formation, arrow: start cocoon assembly, arrow head: BCV membrane rupture. (related to Figure 6)

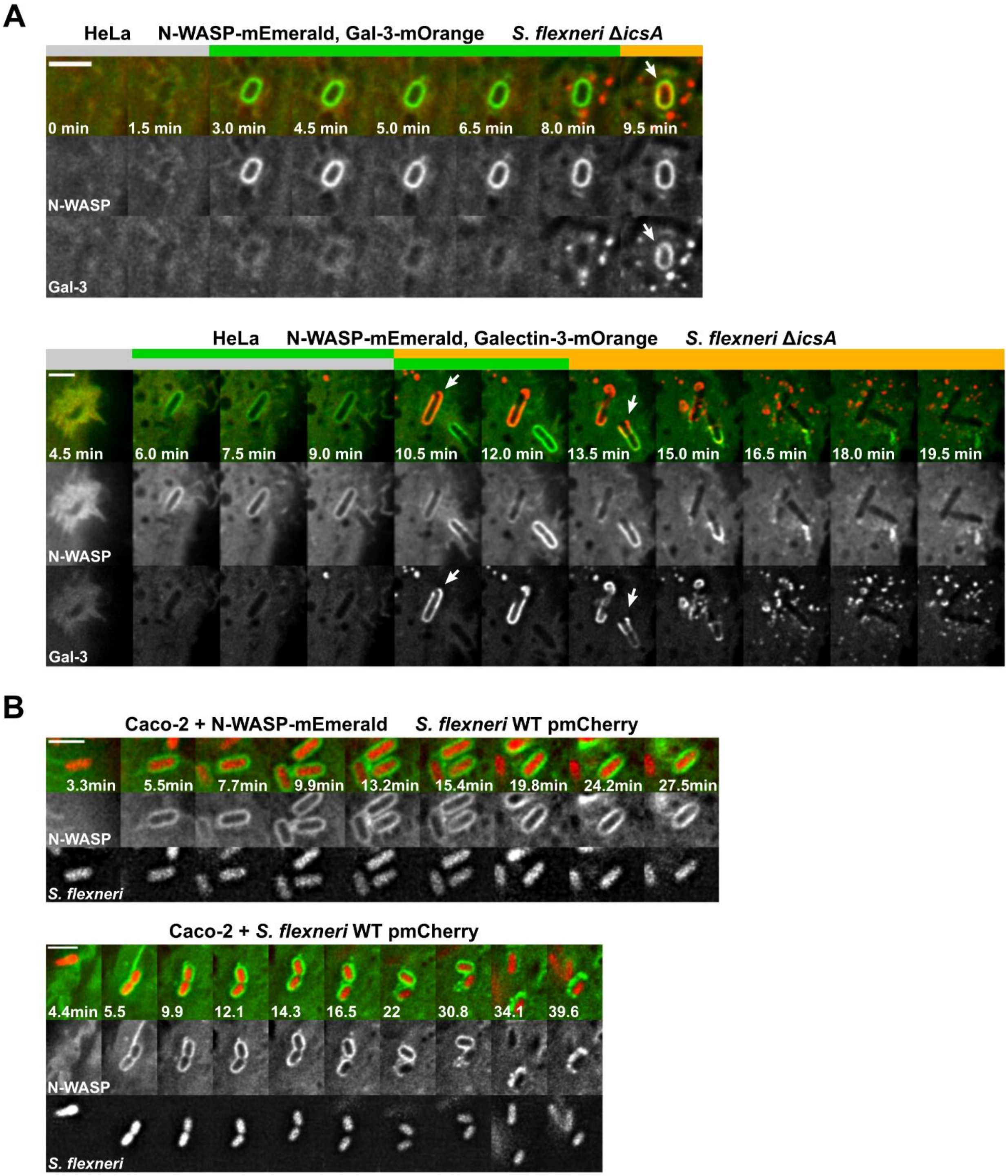


**Figure S7: N-WASP recruitment to the BCV membrane before vacuolar rupture is IcsA-independent**.

(**A**) Two representative time-lapses of HeLa cells co-transfected with N-WASP and galectin-3 and infected with *Shigella* Δ*icsA*. Initial N-WASP recruitment and its constant localization are similar to WT infections (Figure 7). (**B**) Depicted are two time-lapses of mCherry-expressing *S. flexneri* WT (red) invading Caco-2 cells expressing N-WASP. N-WASP is likewise constantly localized around the *Shigella* BCV (scale bars: 3 μm). (related to Figure 6)

#### SUPPLEMENTAL EXPERIMENTAL PROCEDURES

##### Quantitative image analysis

Since almost all invading wild-type *Shigella* escaped into the host cytoplasm in the duration of the experiment, we focused in our analysis on *Shigella* that successfully invaded the cytosol (galectin-3 recruitment to BCV membrane remnant) as indicator for efficient infection. Time-lapse of *Shigella* WT infecting HeLa cells co-expressing actin-GFP and galectin-3-mOrange allowed correlation and precise timing of actin dynamics with vacuolar rupture. Actin cocoon assembly was defined as the *de novo* formation of an actin-enriched structure around the BCV membrane with clear borders that assembled *de novo* after cellular entry and before vacuolar rupture for a duration of >2 min. The comparison of the overall percentage of bacteria that assemble an actin cocoon before vacuolar rupture did represent a solid readout to investigate e.g. the effect of host protein inhibitors or *Shigella* mutant strains. Cases in which cocoons could not be clearly distinguished from surrounding membrane ruffles or IAMs were excluded from analysis (0-2% per experiment, within the error range). The presence or absence of actin cocoons was investigated. Changes in dynamics, shape, or relative fluorescence intensities were not taken into account. This study focused on primary infections and early invasion steps. Secondary infections usually lack the massive membrane ruffling of primary infections and were not taken into account.

The vacuolar rupture time was defined as time span between the onset of initial membrane ruffling and the first appearance of galectin-3 recruitment to the BCV membrane remnant, including successive invasion steps of cellular uptake, actin cocoon formation, maintenance as well as disassembly, and cytosolic escape (Figure S2). The rupture time can only in combination with the actin cocoon readout indicate disturbed cocoon dynamics. Since other endocytic compartments than the BCV (e.g. *E. coli* InvA phagosomes, IAMs) were never galectin-3-positive, its recruitment was used as marker for *Shigella* entering the cytosol. All rupture events were considered including statistic outliers. Significance was likewise confirmed for the geometrical mean and data sets excluding statistical outliers. In screens with varying infection efficiencies (e.g. *Shigella* mutants), only the first invader per cell was taken into account.

To compare the actin cocoon with cellular or pathogen-induced actin structures (Figure 2A), we normalized the effect of extensive membrane ruffling at the bacterial entry site and of varying actin-GFP expression levels in different cells. We measured the fluorescence intensity of on average the 10 most intense cellular actin stress fibers (mainly ventral and dorsal stress fibers, not in close proximity to focal adhesions) of each infected HeLa cell. The average stress fiber intensity was used to normalize the fluorescence intensity of each individual actin structure measurement. In addition, each single measurement was corrected for the cytosolic actin-GFP signal in its immediate vicinity. *E. coli* InvA is a model for canonical phagocytosis with actin assembly and disassembly cycles around phagosomes (actin flashing). The related enteropathogenic *Salmonella* (*S.* Typhimurium) injects some T3SS effector protein homologues to *Shigella*. Data obtained for *E. coli* InvA and *Salmonella* infections were analyzed equally. The maximum intensity of actin assembly was considered. The lifetime of actin coats around *Shigella*'s BCV or *E. coli*'s phagosome (Fig. 2C) was measured as the time interval between initial actin assembly and its complete disassembly. For *E. coli* InvA phagosomes, one cycle of complete actin assembly and disassembly was considered.

All siRNA experiments were performed in parallel with scramble siRNA controls and HeLa cell controls. Septin-7 is an essential septin filament component and its knockdown by RNA interference efficiently inhibits septin filament assembly (Sirianni et al., 2016). Knockdown of ArpC3 was shown previously to be involved in *Shigella* invasion (Mellouk et al., 2014). For Cdc42 siRNA-mediated knockdown or overexpression of Cdc42 mutants, in parallel to the total population we also analyzed the behavior of only the first invading bacterium per cell (Figure S5D,E). Both analyses gave consistent results.

The intensity of host protein recruitment (Figure 6) was quantified using Fiji. To investigate levels of Cdc42 recruitment during *Shigella* infection (Figure 6B), the mean fluorescence intensity of a ROI comprising either only the plasma membrane ruffle that formed the endocytic compartment or the BCV membrane was measured. All values were normalized to the initial signal of Cdc42 recruited to the plasma membrane. We also analyzed the IcsB-dependent localization of Cdc42 and N-WASP around vacuolar

*Shigella* (Figure 6D,H). Both proteins were considered constantly localized if they, after initial recruitment during early infection steps, remained associated with the galectin-3-stained membrane remnant after vacuolar rupture. Proteins were defined as depleted in case they were initially recruited to the intact BCV, but disappeared before vacuolar rupture. Cdc42 and N-WASP were specified as not recruited, if no localization at the vacuolar membrane was observed after scission of newly formed phagosomes. This does not exclude e.g. initial Cdc42 recruitment to the plasma membrane during membrane ruffling.
